# Intrafilament nucleotide exchange in a prokaryotic actin homolog

**DOI:** 10.64898/2026.07.24.740542

**Authors:** Ingrid E. Adriaans, Cyrille Billaudeau, Charlène Cornilleau, Kee Siang Lim, Céline Dinet, Lars D. Renner, Caroline Peron-Cane, Antoine Jégou, Richard W. Wong, Arnaud Chastanet, Alphée Michelot, Rut Carballido-López

## Abstract

Polymerization and disassembly govern the cellular functions of cytoskeletal proteins. Canonical nucleotide-dependent polymers, including actin and microtubules, renew their nucleotide state through subunit subunit dissociation, nucleotide exchange in solution and repolymerization. Yet the assembly dynamics of the membrane-bound bacterial actin MreB remain unresolved. Here, using total internal reflection fluorescence microscopy and high-speed atomic force microscopy, we visualise Bacillus subtilis MreB assembly on supported lipid bilayers in real time. ATP binding drives polymerization into symmetrically elongating filaments, whereas ATP hydrolysis within filaments promotes disassembly. Unexpectedly, nucleotides continuously exchange within membrane-bound filaments without detectable subunit turnover, coupling filament stability to the surrounding nucleotide pool: ATP exchange stabilizes filaments, whereas ADP exchange triggers rapid fragmentation and disassembly. Monte-Carlo modelling and single-cell in vivo imaging support this mechanism. Thus, intrafilament nucleotide exchange enables MreB filaments to renew their nucleotide state without polymer turnover, revealing nucleotide-state rejuvenation as a new mode of biological polymer regulation.

## Main Text

Bacteria, eukaryotic cells and archaea possess actin homologs with conserved structural features and some related functionality, suggesting that the actin gene originated in the common ancestor of all living organisms ^1,2^. MreB proteins are actin homologs essential for viability and cell shape determination in bacteria, and are also present in some archaea ^3,4^. *In vivo*, MreB assembles into membrane-associated short filaments that move processively around the cell circumference and are thought to act as platforms that restrict the diffusion of cell wall synthases and orient their motion circumferentially, promoting cylindrical elongation ^5–7^.

Biological polymers need to assemble in a controlled and reversible manner to fulfil their cellular functions with proper spatial and temporal coordination ^1^. Eukaryotic actins polymerize spontaneously in solution into polar, helical filaments composed of double parallel protofilaments. Actin filament disassembly is controlled by the hydrolysis of the bound nucleotide adenosine triphosphate (ATP) within filament subunits, and a specific subset of binding partners known as actin-binding proteins (ABPs). Release of hydrolyzed γ-phosphate promotes depolymerization of ADP-actin monomers that rapidly exchange nucleotide back to ATP, allowing multiple polymerization cycles ^2^. Within assembled filaments, nucleotides remain effectively trapped in each subunit until depolymerization, preventing any nucleotide exchange ^8^.

In contrast to actin, MreB proteins assemble into straight pairs of antiparallel protofilaments on a lipid membrane surface in the presence of ATP ^9–11^. MreB polymeric assemblies appear stable and long-lived *in vivo* ^5,12^. However, their ultrastructure and the assembly-disassembly dynamics of single MreB double filaments remain unknown. While the ease of eukaryotic actin purification has enabled decades of mechanistic studies on filament assembly dynamics and regulation ^2^, the fact that bacterial MreB proteins bind directly to membranes via hydrophobic sequences that promote protein aggregation has rendered their purification and labelling in a soluble form notoriously difficult, impeding comparable *in vitro* analyses. In particular, how MreB integrates nucleotide-level regulation with membrane interactions to control polymerization remains poorly defined. To characterize the molecular mechanisms governing MreB assembly dynamics, we used state of the art high-resolution imaging and quantitative biophysical approaches to study polymerization of MreB from the model bacterium *Bacillus subtilis* on supported lipid bilayers (SLBs). We show that MreB polymerization is an ATP-binding-driven lipid membrane-templated mechanism. MreB filaments are kinetically symmetrical and their stability on the membrane is regulated by ATP hydrolysis and intrafilament nucleotide exchange. A stochastic Monte-Carlo model and single-cell *in vivo* imaging data support our findings. Nucleotide cycling within intact filaments defines a new class of biopolymer dynamics.

### Polymerization of *Bacillus subtilis* MreB requires ATP and a cardiolipin-containing membrane

Full-length recombinant MreB from *B. subtilis* (hereafter called ‘MreB’), both untagged and fused to a color-tunable variant of the chemogenetic **p**romiscuous **F**luorescence-**A**ctivating and absorption **S**hifting **T**ag (pFAST) ^13^, forms straight pairs of protofilaments on a monolayer of *B. subtilis* lipid extract in the presence of ATP (Fig. 1A-C), as observed by transmission electron microscopy (TEM) of negatively stained samples. Throughout this work, a ‘filament’ refers to one doublet. The pFAST-MreB fusion is fully functional *in vivo* (Fig. S1). At the concentration of MreB used in these experiments (1.5 µM), comparable to the previously reported critical concentration for polymerization (1-2 µM) ^10,14–17^, no filaments were observed in the presence of either ADP or the non-hydrolysable ATP analogs AMPPNP or AMPPCP, or in the absence of lipids (Fig. 1C and Fig. S2A). Divalent cation pre-binding, typically Mg^2+^, is essential for nucleotide binding in both actin and MreB proteins ^9,17–20^. We quantified the binding kinetics of ATP, ADP and AMPPNP to soluble MreB monomers in the presence of Mg^2+^ using fluorescence anisotropy ^8,17^ (Fig. 1D, Fig. S2B-F and Table 1). The ATP and ADP binding rates were markedly slow, ∼300-fold lower than those reported for actin (Table 1) and ∼10–20-fold lower than those of MreB from *Geobacillus stearothermophilus* ^17^, with anisotropy increases occurring over extended periods compared to diffusion-limited binding (Fig. S2C, D). ATP and ADP exhibited similarly low association rates, suggesting a shared rate-limiting step. A plausible explanation is that MreB, which was purified in a nucleotide-free state, is partially unfolded in the absence of nucleotide. Nucleotide binding may require conformational rearrangements to reach a binding-competent state, slowing association. Chase experiments confirmed that the anisotropy signal arises from nucleotide–protein interactions rather than nonspecific effects (Fig. S2C, D). The equilibrium dissociation constants for both ATP and ADP were in the micromolar range, while they are in the nanomolar range for monomeric actin (Table 1). This indicates moderate affinity of MreB for either ATP or ADP, in contrast to the very high affinity of actin for the two nucleotides. Furthermore, both ATP and ADP were found to bind monomeric MreB with similar affinities, as recently reported for *Geobacillus* MreB ^17^, while Mg^2+^-actin has a 4-fold higher affinity for ATP than for ADP (Table 1). No binding was detected for AMPPNP (Fig. 1D, S2B), explaining the absence of polymers in the presence of this ATP analog (Fig. S2A).

**Fig. 1.**
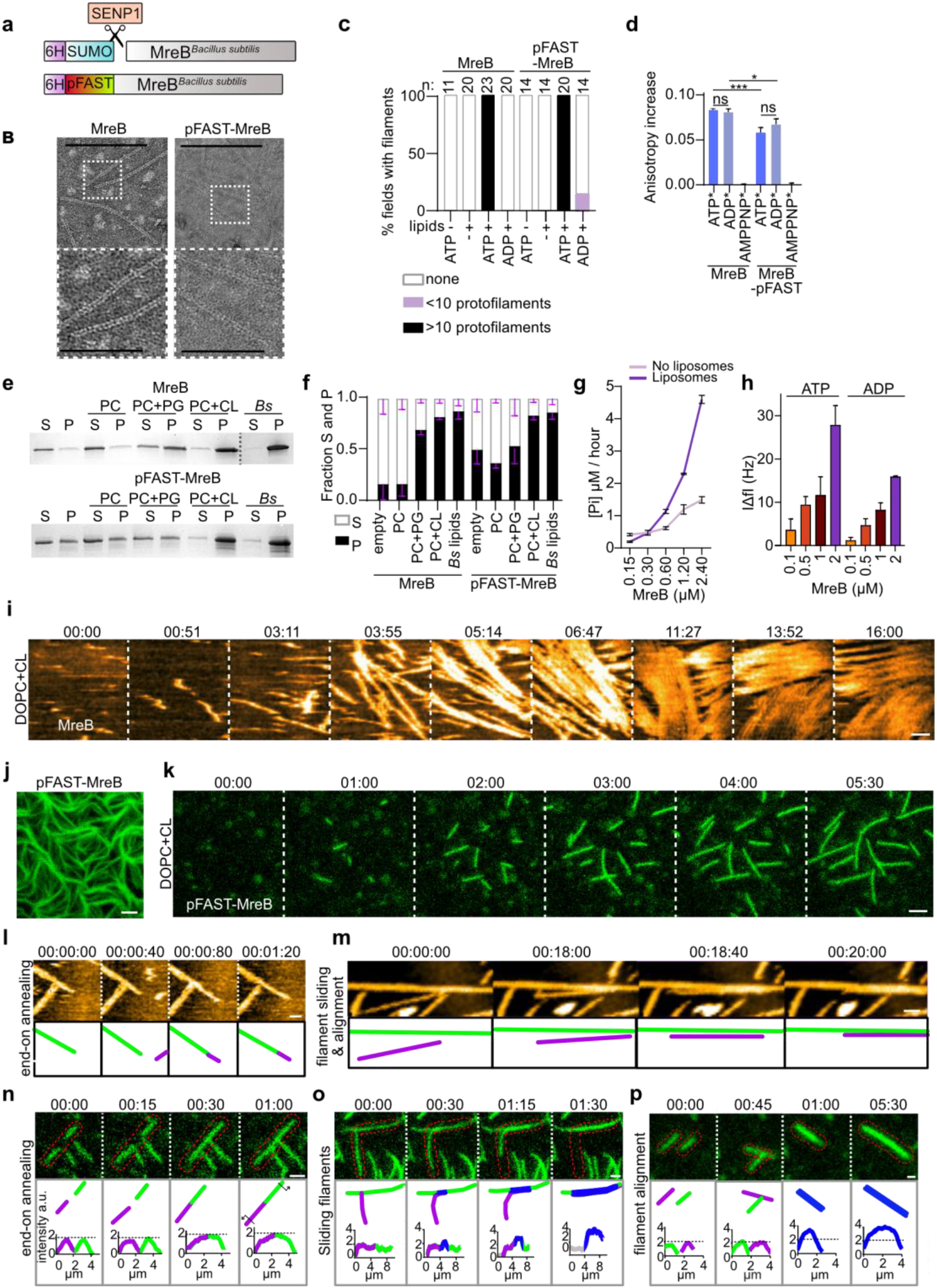
Polymerization of *B. subtilis* MreB requires ATP and anionic phospholipids. (A) Scheme of the two *B. subtilis* MreB constructs used in this study, namely untagged MreB obtained by cleaving of 6xHis (6H)-SUMO by the protease SENP1 (‘MreB’), and 6xHis-pFAST-tagged MreB. (‘pFAST-MreB’). (B) Representative TEM images of pairs of protofilaments formed by tagged and untagged MreB (1.5 µM) on a lipid monolayer of *B. subtilis* lipid extract in the presence of ATP (100 µM). Scale bars, 200 nm and 50 nm in the enlargement. (C) Quantification of the presence of MreB pairs of filaments on TEM fields in the presence or absence of nucleotide (100 µM) and lipids. n, number of TEM fields analyzed. (D) Steady-state fluorescent anisotropy values of N^6^(6-Aminohexyl)-ATP-Atto-488 (ATP*), N^6^-(6-Aminohexyl)-ADP-ATTO-488 (ADP*) and N^6^-(6-Aminohexyl)-AMPPNP-ATTO-488 (AMPPNP*) (0.2 µM) binding to MreB and pFAST-MreB (1 µM). Bars represent mean ± standard deviation. Values are representative of three independent experiments. (E) Representative Coomassie-stained 4-15% SDS-PAGE gel of liposome binding assay of MreB (2 µM) in the presence of ATP (200 µM). S, supernatant; P, pellet; PC, 1,2-dioleoyl-sn-glycero-3-phosphocholine (DOPC); PG, phosphatidylglycerol; CL, cardiolipin; Bs, *B. subtilis* total lipid extract. Ratio (Mol%) PC:PG and PC:CL is 80:20. In the absence of binding, MreB remained in the supernatant, whereas liposome-associated MreB was recovered in the pellet. Control reactions without liposomes were included to account for nonspecific MreB precipitation. (F) Quantification of 3 independent liposome binding assays. (G) ATPase activity, measured by monitoring inorganic phosphate (P_i_) release, of MreB at the indicated concentrations in the presence of 0.5 mM ATP and in the presence or absence of 0.5 mg/mL liposomes. Values are means with standard deviation. Results of two independent experiments. (H) Adsorption of MreB to a DOPC:CL (80:20) SLB in the presence of ATP or ADP (2 mM) measured by QCM-D. Bars represent mean ± standard deviation of four independent experiments for ATP and two independent experiments for ADP and AMPPNP. (I) HS-AFM of 1.2 µM untagged MreB on a SLB (DOPC: CL 80:20) in the presence of 0.2 mM ATP. Field of view, 1.2×1.2 µm. Scan rate, 1 frame/0.4 sec. Time in minutes:seconds. Scale bar, 200 nm. (J, K) TIRF microscopy of 1.2 µM (J) and 0.05 µM (K) pFAST-MreB on a SLB (DOPC:CL 80:20) in the presence of 0.2 mM ATP. Field of view, 29.44×26.18 µm. Time in minutes:seconds. Scale bar, 2 µm. (L-P) Examples of end-to-end filament annealing (L, N) and of filament lateral alignment (M, P) and sliding (M, O) visualized by HS-AFM (0.6 µM MreB. Time in min:sec:hundredths of second. Scale bars, 100 nm) (L,M) or TIRF (0.05 µM MreB. Time in min:sec. Scale bars, 1 µm) (N-P). Cartoons below each panel depict individual filaments in different couleurs for visual help. Plots in the TIRF panels (N-P) show fluorescence intensity profiles along the regions outlined in red in the corresponding TIRF images. Individual filaments are shown in green and purple; laterally aligned filaments in blue, no filament in grey.

**Table 1.** Biochemical parameters: Actin (ACTA2) versus MreB^Bs^.

| Parameter | Name | Mg-Actin ACTA2 [refs.] | Mg-MreB<sup>Bs</sup> [this study] |
| --- | --- | --- | --- |
| ATP binding rate to monomers | $k_{+,G,ATP}$ | $3.2 \mu\text{M}^{-1} \cdot \text{s}^{-1}$ <sup>40</sup> | $0.01 \mu\text{M}^{-1} \cdot \text{s}^{-1}$ ($0.05 \mu\text{M}^{-1} \cdot \text{s}^{-1}$ for E136A) (ND for D158A) |
| ATP dissociation rate from monomers | $k_{-,G,ATP}$ | $0.0012 - 0.0045 \text{ s}^{-1}$ <sup>8,36,41</sup> | $0.005 \text{ s}^{-1}$ ($0.004 \text{ s}^{-1}$ for E136A) (ND for D158A) |
| ADP binding rate to monomers | $k_{+,G,ADP}$ | ND (fast - considered similar to $k_{+,G,ATP}$) | $0.013 \mu\text{M}^{-1} \cdot \text{s}^{-1}$ ($0.05 \mu\text{M}^{-1} \cdot \text{s}^{-1}$ for E136A) (ND for D158A) |
| ADP dissociation rate from monomers | $k_{-,G,ADP}$ | $0.01 - 0.09 \text{ s}^{-1}$ <sup>8,36,41</sup> | $0.0073 \text{ s}^{-1}$ ($0.008 \text{ s}^{-1}$ for E136A) (ND for D158A) |
| ATP dissociation constant | $K_{d,ATP,G}$ | $0.375\text{-}1.40 \text{ nM}$ | $0.50 \mu\text{M}$ ($0.08 \mu\text{M}$ for E136A) (ND for D158A) |
| ADP dissociation constant | $K_{d,ADP,G}$ | $3.12\text{-}28.12 \text{ nM}$ | $0.56 \mu\text{M}$ ($0.16 \mu\text{M}$ for E135A) (ND for E158A) |
| Affinity of monomers for ADP, relative to ATP | $\frac{K_{d,ADP,G}}{K_{d,ATP,G}}$ | $\approx 4$ <sup>8,36,41</sup> | $\approx 1$ |
| CC of assembly for ATP-monomers | $c_{c,ATP}$ | $0.15 \mu\text{M}$ <sup>42,43</sup> | negligible ($0.003 \mu\text{M}$ for WT, E136A and D158A) |
| CC of assembly for ADP-monomers | $c_{c,ADP}$ | $1.2 \mu\text{M}$ <sup>36</sup> | $> 1.5 \mu\text{M}$ (if possible) |
| ATP-monomer AR constant at BE | $k_{+,b,t}$ | $11.6 \mu\text{M}^{-1} \cdot \text{s}^{-1}$ <sup>25</sup> | $45 \mu\text{M}^{-1} \cdot \text{s}^{-1}$<br>(for WT, E136A and D158A) |
| ATP-monomer AR constant at PE | $k_{+,p,t}$ | $1.3 \mu\text{M}^{-1} \cdot \text{s}^{-1}$ <sup>25</sup> | |
| ADP-monomer AR constant at BE | $k_{+,b,d}$ | $3.8 \mu\text{M}^{-1} \cdot \text{s}^{-1}$ <sup>25</sup> | ND |
| ADP-monomer AR constant at PE | $k_{+,p,d}$ | $0.16 \mu\text{M}^{-1} \cdot \text{s}^{-1}$ <sup>25</sup> | |
| ATP-subunit DR constant at BE | $k_{-,b,t}$ | $1.4 \text{ s}^{-1}$ <sup>25</sup> | low ($0.135 \text{ s}^{-1}$ for WT, E136A and D158A) |
| ATP-subunit DR constant at PE | $k_{-,p,t}$ | $0.8 \text{ s}^{-1}$ <sup>25,44</sup> | |
| ADP-Pi-subunit DR constant at BE | $k_{-,b,dp}$ | $0.16 \pm 0.07 \text{ s}^{-1}$ <sup>35</sup><br>$0.33 \pm 0.16 \text{ s}^{-1}$ <sup>35</sup><br>$1.5 \pm 0.4 \text{ s}^{-1}$ <sup>35</sup><br>$0.6 \text{ s}^{-1}$ <sup>45</sup> | ND |
| ADP-Pi-subunit DR constant at PE | $k_{-,p,dp}$ | $0.15 \text{ s}^{-1}$ <sup>45</sup><br>$0.16 \pm 0.07 \text{ s}^{-1}$ <sup>35</sup> | |
| ADP-subunit DR constant at BE | $k_{-,b,d}$ | $7.2 \text{ s}^{-1}$ <sup>25</sup><br>$5.8 \pm 0.4 \text{ s}^{-1}$ <sup>35</sup> | $60 \text{ s}^{-1}$ |
| ADP-subunit DR constant at PE | $k_{-,p,d}$ | $0.27 \text{ s}^{-1}$ <sup>25</sup><br>$0.14 \pm 0.04 \text{ s}^{-1}$ <sup>44</sup> | |
| nucleotide-free-subunit DR constant at BE | $k_{-,b,e}$ | ND | $4 \text{ s}^{-1}$ |
| nucleotide-free-subunit DR constant at PE | $k_{-,p,e}$ | ND | |
| ATP hydrolysis rate in filaments | $k_{hyd}$ | $0.3 \text{ s}^{-1}$ <sup>46</sup> | $5 \cdot 10^{-4} \text{ s}^{-1}$ (with liposomes)<br>$2.9 \cdot 10^{-4} \text{ s}^{-1}$ (without liposomes)<br>($\approx 0 \text{ s}^{-1}$ for E136A and D158A)<br>(combined rate hydrolysis + Pi release) |
| Pi release rate in filaments | $k_{pir}$ | $0.0068 \pm 0.0021 \text{ s}^{-1}$ <sup>35</sup><br>$0.002 - 0.006 \text{ s}^{-1}$ <sup>45,47</sup> | |
| ADP-to-ATP nucleotide exchange at the filament side | $k_{ex,F}$ | $0 \text{ s}^{-1}$ <sup>8</sup> | $0.014 \text{ s}^{-1}$ ($\approx 0$ for E136A and ND for D158A) |

BE, barbed end; PE, pointed end; AR, association rate; DR, dissociation rate; CC, Critical concentration

We then set out to analyze MreB assembly dynamics on supported lipid bilayers (SLBs), which mimic the native, fluid environment of the plasma membrane. Bacterial lipid extracts contain lipids with different charges and geometries, making it difficult to form flat, homogeneous SLBs suitable for surface-sensitive, high-resolution microscopy studies ^21,22^. Anchoring of MreB filaments to a lipid surface is mediated by short hydrophobic sequences that protrude from the monomers and insert into the lipid bilayer, but electrostatic interactions with anionic phospholipids are required for initial lipid binding ^10,11,17^. Through liposome co-sedimentation assays, we found that doping 1,2-dioleoyl-sn-glycero-3-phosphocholine (DOPC) with negatively-charged cardiolipin (CL), and phosphatidylglycerol (PG) to a lesser extent, promote MreB membrane association in the presence of ATP (Fig. 1E, F). This is consistent with recent molecular dynamics simulations showing that *Escherichia coli* MreB filaments recruit CL ^18^. CL is a cone-shaped, tetra-acylated anionic phospholipid composed of two phosphatidyl moieties linked by glycerol. Despite its structural similarity to two cylindrical-shaped PG molecules, its net charge equals that of a single PG at neutral pH, suggesting that the preferential binding to CL over PG may be related to geometric cues. Binding of MreB to liposomes composed of *B. subtilis* lipid extract or containing 20% CL (similar to the proportion of CL in lipid extracts from growing *B. subtilis* cells ^23,24^) was comparable (Fig. 1E, F), suggesting that CL could function as a major determinant of MreB membrane association in *B. subtilis*. Release of inorganic phosphate (Pi) from MreB after ATP hydrolysis was increased in the presence of liposomes (Fig. 1G) ^10^, indicating that MreB ATPase activity is induced upon or after membrane binding.

SLBs containing 20% CL formed flat, smooth surfaces with fluid lipid diffusion, as shown by fluorescence recovery after photobleaching (FRAP) (Fig. S3A, B, Movie 1). MreB bound to the SLB in the presence of either ATP or ADP, although ATP significantly enhanced membrane binding (Fig. 1H and Fig. S4A, B). We next visualized the polymerization dynamics of 1.2 µM untagged MreB filaments on MICA-supported 20% CL SLBs by high-speed atomic force microscopy (HS-AFM) (Fig. 1I, Movie 2). Filaments rapidly appeared in the scan area, elongated and increased in density, while displaying high lateral mobility on the fluid lipid bilayer, mostly due to displacement by the AFM tip during scanning. Most filaments grew beyond the field of view (1.2 × 1.2 µm) and, at high densities, aligned laterally displaying nematic order (Fig.1I). To follow MreB filament elongation, we then monitored the assembly of pFAST-MreB on a larger field of view using total internal reflection fluorescence (TIRF) microscopy and the fluorogenic pFAST ligand TFLime (green) (Fig. 1J, K). At 1.2 µM MreB, polymerization also resulted in a very dense nematic network of micrometer-long filaments (Fig. 1J). We lowered the MreB concentration to decrease filament density in order to visualize individual filaments. At 0.05 µM MreB, isolated short filaments appeared within one minute of incubation and progressively elongated (Fig. 1K, Movie 3). Filaments were slightly bent, with a median radius of curvature of 1/0.11 µm (and 1/0.22 µm at 1.2 µM MreB) (Fig. S3C). End-to-end annealing was also observed, which did not interrupt filament growth (Fig. 1L and N). When filaments came into contact, they changed orientation and aligned laterally, and often slid along one another (Fig.1M, O and P).

### MreB filaments elongation is kinetically symmetrical

High resolution HS-AFM images confirmed that we were imaging single MreB doublets (Fig 2A and Fig. S3D). In TIRF microscopy images, the mean fluorescence intensity of filaments revealed a dominant population of uniform intensity, and additional minor populations of two-and three-fold higher intensities, consistent with bundles of double and triple filaments, respectively (Fig. S3E). Next, we attempted to quantify the polymerization dynamics of individual filaments of untagged MreB. Due to their high lateral mobility, individual filaments frequently traversed the limited HS-AFM scan area before stable tracking could be established. Consequently, our quantitative HS-AFM analysis (Fig. 2B, C) was restricted to sub-micrometer filament segments exhibiting transient immobilization or reduced mobility, likely due to transient interactions with the AFM tip. Because of these tracking difficulties, only a very limited number of filaments could be tracked for a short timeframe and thus the calculated HS-AFM elongation rate (14.4 ± 10.16 nm·s⁻¹, n=7) serves as first estimate of assembly (Fig. 2B-D, Movie 4).

**Fig. 2.**
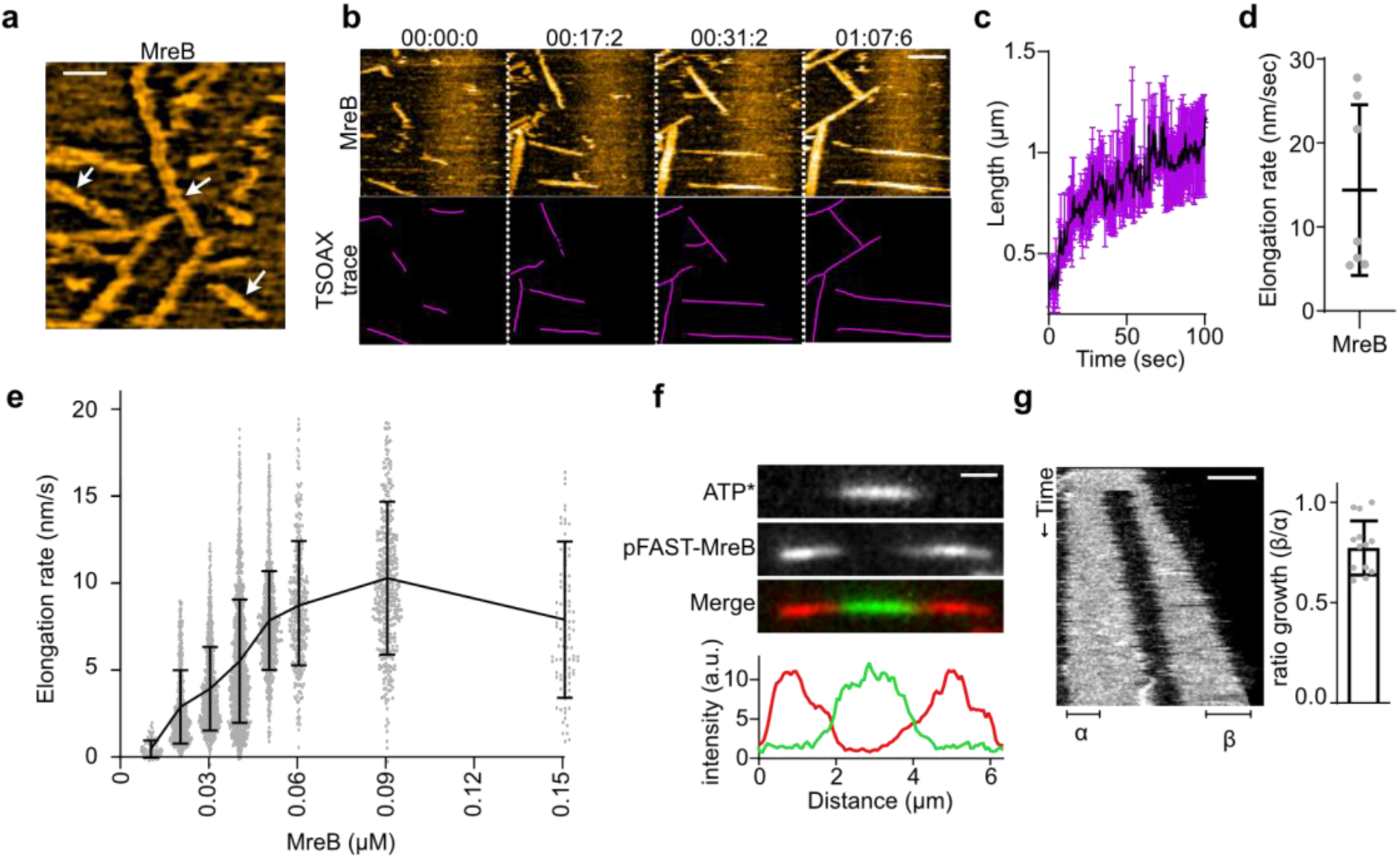
MreB pairs of filaments elongate symmetrically. (A) High-resolution, Fast Fourier Transform (FFT) bandpass filtered HS-AFM image showing MreB doublets (arrows). Scale bar, 100 nm. (B) Representative HS-AFM time-lapse of MreB (0.6 µM) assembly on a DOPC:CL (80:20) SLB. The bottom panel shows the TSOAX traces used to track lengths and calculate elongation rate. Time in min:sec:hundredths of second. Scale bar, 300 nm. (C, D) average MreB filament length over time (C) and corresponding elongation rate (D) in HS-AFM experiments. Bars represent mean ± standard deviation. n = 7 from two independent experiments. (E) Filament elongation rate relative to pFAST-MreB concentration in TIRF experiments. n = 77, 636, 849, 1142, 674, 348, 357 and 105 for 0.01, 0.02, 0.03, 0.04, 0.05, 0,06, 0.09 and 0.15 µM MreB, respectively. (F) *Top panel,* MreB filament first labelled by fluorescent ATP (N^6^-(6-Aminohexyl)-ATP-ATTO-488) (ATP*, green) before addition of pFAST-MreB labelled with TFCoral (red). Scale bar, 1 µm. *Bottom panel,* corresponding fluorescence intensity along the filament in the green and red channels. (G) *Left panel*, representative kymograph of a photobleaching experiment on a single pFAST-MreB filament labelled with TFLime. Vertical axis, time (2 sec/frame). Scale bar, 2 µm. *Right panel*, ratio of filament growth on both filament ends after photobleaching. Growth at both filament ends (designated α and β) was quantified, and the ratio of the shorter to the longer growth length was calculated to give a ratio of the growth, where a ratio approaching 0 indicates polar growth restricted to a single filament end, and ratios close to 1 indicate non-polar, identical growth at both ends. By defining the ratio using the shorter and longer extension lengths, filaments were effectively assigned an artificial orientation during the analysis. As a result, stochastic fluctuations in subunit addition between the two ends, even in the absence of intrinsic polarity, would generate a small apparent asymmetry in growth. n = 13.

Filament elongation could, however, be followed over several micrometers by TIRF microscopy using pFAST-MreB labelled with TFLime (Fig. S3F, G, Movie 5). Elongation rates increased linearly with MreB concentration at low MreB concentrations, and reached a plateau above 0.06-0.09 µM MreB (Fig. 2E). A linear fit at low concentrations yielded the association rate constant*k*_+,*t*_ ≈ 45 µ*M*. *s*^−1^, the dissociation rate constant *k*_−,*t*_ ≈ 0.135 *s*^−1^, and the critical concentration *C_c_* ≈ 0.003 µ*M* (Table 1, see Supporting Data for calculations details) ^25,26^. Notably, the elongation rate value at the plateau was consistent with the HS-AFM-derived rate obtained at much higher protein concentration (1.2 µM), suggesting that the plateau observed in TIRF experiments (Fig. 2E) is not an artefact of the pFAST tag or its labelling strategy. Such plateau contrasts with the polymerization of actin filaments, which displays a linear dependence on monomer concentration well beyond the micromolar range. A likely explanation is that actin polymerization relies on the rapid 3D-diffusion of actin monomers in solution (estimated by the Stokes-Einstein at 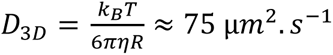, with *η* = 1 *mPa*. *s* and *R* = 3 *nm*), whereas MreB polymerization may first require membrane association of subunits followed by slower 2D-diffusion on the lipid surface (lipid-anchored protein typically exhibit lateral diffusions *D*_2*D*_ < 2 µ*m*^2^. *s*^−1^ ^27^). Membrane-bound subunits at filament ends might also transiently dissociate, undergo slow membrane diffusion and rebind with high efficiency.

Taken together, these findings suggest a model in which ATP-MreB subunit addition to the filament end is predominantly/exclusively mediated by membrane-bound subunits. Membrane recruitment may increase the local concentration of ATP-MreB and/or induce a conformational change that renders it polymerization-competent, enabling efficient incorporation at filaments ends, with subunit association greatly exceeding dissociation.

We next compared polymerization dynamics at both ends of filaments in dual-color TIRF experiments. While actin filaments are structurally polar, crystal structures and *in vivo* crosslinking suggested that MreB might form pairs of antiparallel protofilaments, implying that they display non-polar growth ^9^. We polymerized MreB in the presence of ATP mixed with the fluorescent ATP analog N^6^(6-Aminohexyl)-ATP-ATTO-488 (ATP*, green) (17:1), and then added pFAST-MreB labelled with TFCoral (red). This low fraction of ATP*-labelled monomers reduces filament density and polymerization rates by two-fold, possibly owing to steric conflicts with ATP*, but it does not affect MreB filaments structure or length (Fig. S5), indicating that ATP* is less efficient for polymerization than ATP but structurally compatible and thus functional to label MreB filaments ^8^. Chase anisotropy measurements confirmed specific binding of ATP* to the ATP-binding pocket of MreB, as the signal was competitively displaced by excess unlabeled ATP (Fig S2C). Upon buffer exchange, newly added red subunits accumulated symmetrically at both ends of preexisting green filaments (Fig. 2F), confirming bidirectional elongation. Independent evidence was obtained by photobleaching a central region of a single filament, and measuring elongation rates at both filament ends, which were found to be comparable (Fig. 2G). One end occasionally elongated slightly faster than the other. This weak apparent polarity likely results from crowding in the SLBs, with membrane or neighboring filaments blocking or obstructing filament ends, and from stochastic fluctuations in subunit addition amplified by the growth-ratio definition used in the analysis (see Fig. 2G legend).

### ATP hydrolysis is not required for MreB polymerization

It was previously suggested that ATP hydrolysis mediates MreB filament formation at the membrane ^10^. Consistent with this, we only detected filaments in the presence of ATP (Fig 1C), and lipids stimulated Pi release (Fig. 1G). To test whether Pi release is required for filament formation and/or stabilization on the membrane, we tested the effect of ADP and the phosphate structural analog Aluminum Fluoride (ADP-AlF_4_^-^). AlF_4_^-^ mimics the phosphoryl group in the catalytic transition state (or “ready to be split” state) after ATP hydrolysis but before Pi dissociation (ADP-Pi state), and has been shown to stabilize ADP-actin filaments ^28^. In the presence of ADP-AlF_4_^-^, MreB formed filaments similar to those observed with ATP, albeit with a slightly slower polymerization rate (Fig. S6A-C), whereas no filaments were observed. in the presence of ADP.

To determine whether ATP hydrolysis itself is actually required for polymerization, we tested the effect of mutations in two highly conserved residues within actin-like proteins that are believed to impair ATP hydrolysis, E136A and D158A. In the prokaryotic actin-like ParM protein, the equivalent E136A mutation abolishes detectable ATPase activity ^29^, whereas the mutation equivalent to D158A markedly reduces ATPase activity, albeit not ATP binding, in actin ^30,31^. The ATPase activities of these MreB mutants have not been characterized *in vitro*, but vegetatively growing *B. subtilis* cells carrying the MreB^E136A^ and MreB^D158A^ mutations display mild defects in cell morphology and MreB localization ^7,29,32,33^. We found that both mutants display markedly reduced Pi release relative to wild-type (WT) MreB (Fig 3A). However, E136A has higher affinity for both ATP and ADP than WT MreB, whereas D158A exhibits reduced ATP affinity and no detectable ADP binding (Fig 3B, Fig S7A-E). We observed similar effects for E136A and D158A of *G. stearothermophilus* MreB (De San Eustaquio-Campillo *et al.,* in preparation). Notably, both mutants assembled into filaments indistinguishable from WT MreB filaments (Fig. 3C and Fig. S7A) with similar kinetics (Fig. 3C, D). Taken together, these results indicated that, as in actin, ATP hydrolysis is not required for MreB polymerization but occurs later, within the filament.

**Fig. 3.**
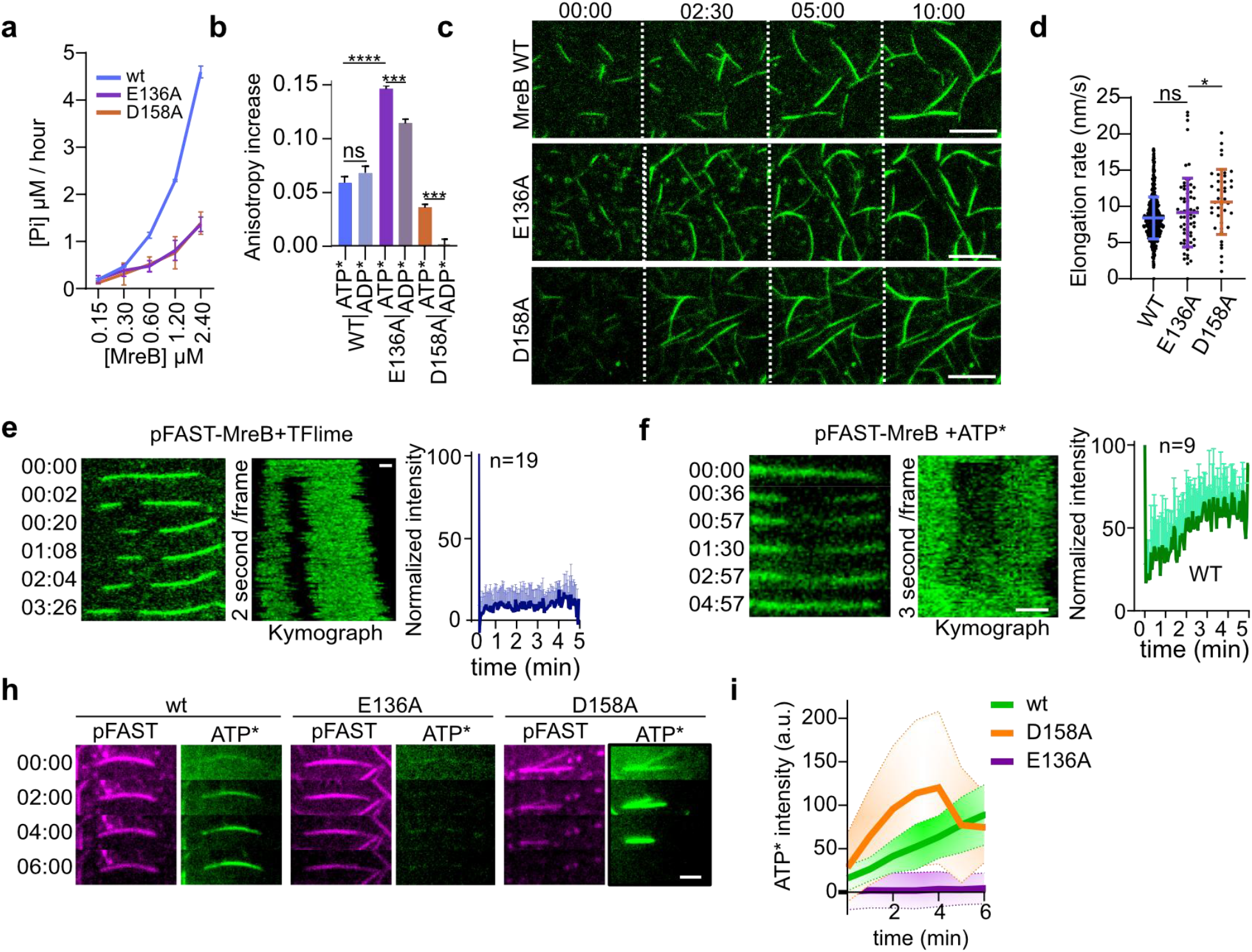
ATP hydrolysis is dispensable for polymerization, and MreB filaments exchange ATP with the solution. (A) Markedly reduced ATPase activity of the E136A and D158A mutants, measured by monitoring inorganic phosphate (Pi) release at the indicated concentrations in the presence of 0.5 mM ATP and in the presence of 0.5 mg/mL liposomes. Values are representative of two independent experiments. (B) Fluorescent anisotropy of N^6^(6-Aminohexyl)-ATP-Atto-488 (ATP*) and N^6^(6-Aminohexyl)-ADP-Atto-488 (ADP*) of mutants E136A and D158A and WT MreB from figure 1D used for comparison. Values are representative of three independent experiments. (C, D) Representative time-lapses (C) and elongation rates (D) of MreB WT and ATPase mutants (0.05 µM) filament growth on DOPC:CL SLB. Time in minutes:seconds. Scale bars, 5 µm. TSOAX was used for measuring filament length over time and determining the elongation rate. n = 318, 68 and 39 for WT, E136A and D158A respectively from two independent experiments. (E,F) FRAP experiments on single filaments of WT MreB (0.05 µM pFAST-MreB) labelled with TFLime (E) or fluorescent ATP* (17:1 ATP:ATP* 50:3 µM) (F). Time in minutes:seconds. Standard deviation bars on top only. Scale bars, 1 µm. *Left panel*, representative time-lapse; *Middle panel*, kymograph; *Right panel*, graph with FRAP result. (H) Time-lapse of incorporation of 3 mM pure fluorescent ATP* (green) along preformed pFAST-MreB filaments (0.05 µM) labelled with TFCoral (magenta). ATP* (green) signal for E136A is enhanced to detect signal. Time in minutes:seconds. Scale bars, 2 µm. (I) Corresponding quantification. MreB WT, green (n=19, initial slope indicates 12.2 intensity min^-1^); E136A, purple (n=12, initial slope indicates 0.5 intensity min^-1^); D158A, orange (n=17, initial slope indicates 28.8 intensity min^-1^). Solid line, mean; dotted lines, standard deviation.

### Nucleotide exchange within membrane-bound MreB filaments

Under limiting ATP concentrations, *Geobacillus* MreB filaments progressively disappeared from TEM fields over time, whereas in the presence of excess ATP, they displayed similar density and lengths for incubation times ranging from minutes to several hours, while continuously releasing P_i_ at a constant rate ^10^. These observations led to the proposal that P_i_ release promotes depolymerization, similar to actin, and that filaments turnover on the membrane. However, in the presence of excess ATP, *B. subtilis* MreB filaments elongated and then remained stable through TIRF time courses (up to several hours), with no detectable depolymerization, and released P_i_ very slowly, at a rate of 0.030 ± 0.001 Pi/min/MreB molecule (∼1 P_i_ released by MreB molecule in 33 min) (Fig. 1G), which is several times slower than the P_i_ release rate reported for actin filaments (Table 1) ^28,34,35^. FRAP experiments of TFLime-labeled filaments showed that monomers do not turnover within filaments (Fig. 3E). Although TFLime binding to pFAST is reversible under normal imaging conditions ^13^, the lack of FRAP recovery suggests that the high-intensity laser illumination used for photobleaching induces irreversible photochemical trapping of the fluorogen on the tag, preventing exchange with unbleached TFLime molecules. However, ATP*-labelled filaments showed rapid signal recovery, indicating nucleotide exchange between the filaments and the solution (Fig. 3F). No significant recovery was detected for the ATPase-deficient mutant E136A (Fig. S7F). Under these standard ATP*-labelling conditions (ATP:ATP* 17:1), D158A filaments were not visible, likely because of the reduced nucleotide binding affinity of this mutant (Fig. 3B and Fig. S7D, E) together with the lower efficiency of ATP* over ATP for polymerization (Fig. S5C, F). When preformed pFAST-MreB filaments labelled with TFCoral were exposed to pure ATP* instead, homogeneous binding of ATP* was observed along the length of both WT and D158A filaments, but not of E136A filaments (Fig. 3H). Under these conditions, D158A filaments incorporated the nucleotide about twice as fast as WT filaments (Fig. 3I).

Eukaryotic actin filaments show no evidence of rapid nucleotide exchange with the solution ^8,36^, consistent with their atomic model ^37^ where the helical twist and constrained interdomain motions within the filament irreversibly block the release of the nucleotide. Crystal structures of MreB protofilaments in solution and molecular dynamic simulations suggest that polymerization closes the nucleotide-binding pocket of MreB to promote nucleotide hydrolysis, causing subunit flattening, protofilament bending and a twist angle ^9,19,38^. However, membrane anchoring was not or only poorly considered in these studies. Polymerization of otherwise intrinsically curved and slightly twisted MreB filaments directly on a membrane surface would reduce the twist angle and straighten the filaments ^9,19,38^. These and/or subsequent ATP hydrolysis could cause some conformational changes that open the nucleotide binding pocket, allowing intrafilament nucleotide exchange. To our knowledge, this is the first report of nucleotide cycling within a biological polymer.

### Free ADP induces depolymerization of MreB filaments

Based on our results, we hypothesized that, in addition to Pi release following ATP hydrolysis promoting dissociation – a mechanism shared with actin – nucleotide exchange along filaments may serve as a complementary mechanism for tuning their nucleotide composition and stability. To allow Pi release to proceed in the absence of ATP cycling, we polymerized MreB in the presence of ATP and then washed off the polymerizing solution and replaced it with MreB-free and nucleotide-free buffer. After a few minutes, WT filaments began to undergo very slow disassembly from both ends (−1.03 ± 0.54 nm.s^-1^), which progressively accelerated until the filaments had vanished from the membrane in ∼20 min (Fig 4A-C). By contrast, D158A filaments depolymerized much faster, within 5 min (−14.67 ± 9.14 nm.s^-1^), whereas E136A filaments remained stable during the length of the experiment (Fig 4A-C).

**Fig. 4.**
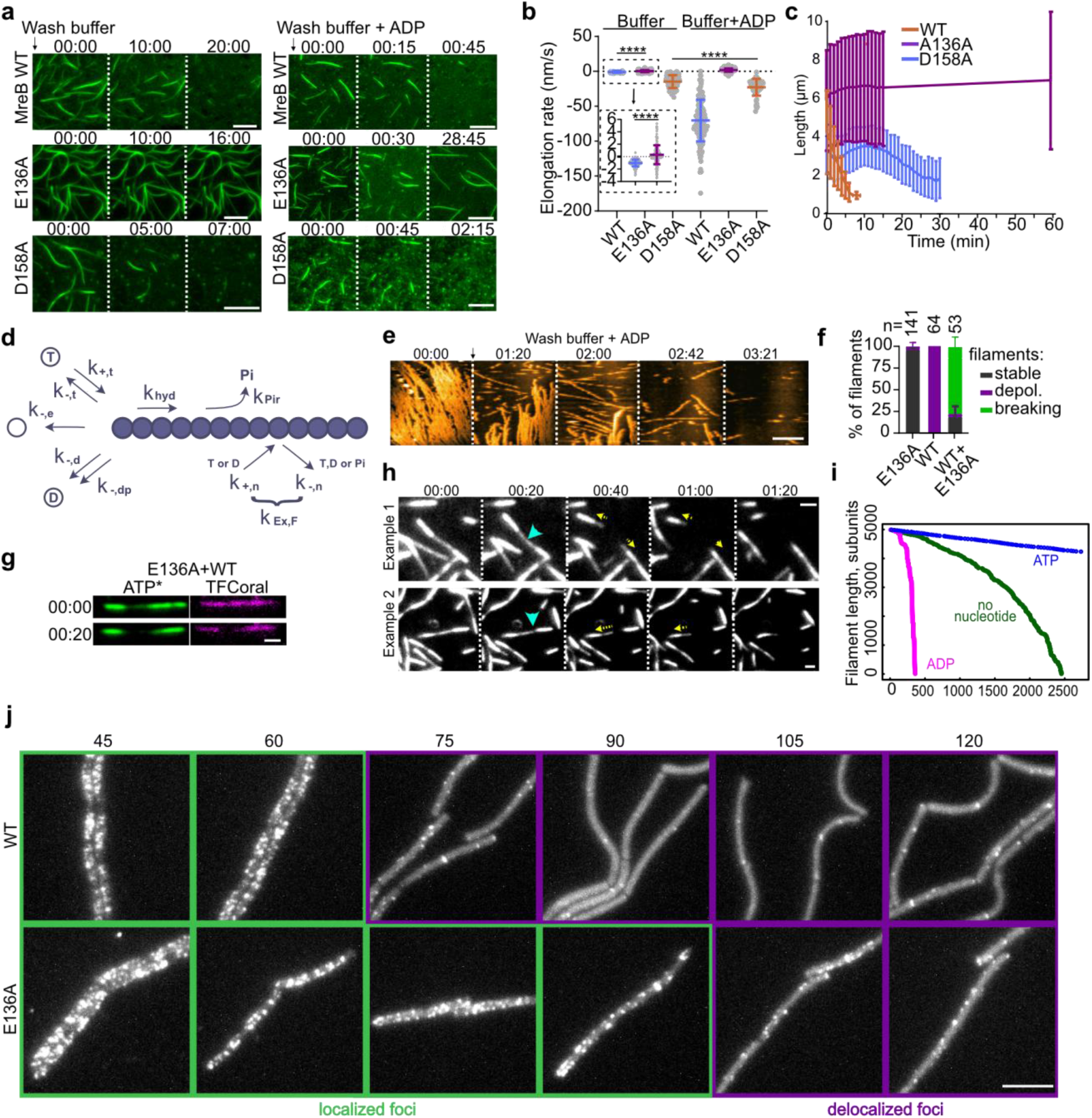
ADP triggers rapid depolymerization of MreB filaments. (A) TIRF microscopy time-course of the effect of replacing the polymerization buffer with MreB-free and nucleotide-free buffer (*Left panels*) or with MreB-free ADP-containing (0.2 mM) buffer (*Right panels*) on pFAST-MreB (0.05 µM) filaments labelled with TFLime on an SLB (DOPC:CL 80:20). Time in minutes:seconds. Scale bars, 5 µm. (B) Quantification of the mean disassembly rates (negative elongation rates) of the experiment shown in A. The square with dotted-line indicates the zoomed area. n = 219, 206, 87, 157, 408 and 60 for WT, E136A, D158A buffer wash and WT, E136A, D158A buffer + ADP wash, respectively. (C) Tracking of filament length over time shows that WT MreB disassembly is not linear and accelerates after 15 min. (D) Schematics of the reactions considered in the model. MreB filament subunits (in blue) can be in any of the four state: ATP-bound, ADP-Pi-bound, ADP-bound, or nucleotide free. Standard values for WT MreB used for the simulations are: *k*_+,*t*_ = 45 µ*M*^−1^*s*^−1^, *k*_−,*t*_ = *k*_−,*dp*_ = 0.135 *s*^−1^, *k*_−,*d*_ = 60 *s*^−1^, *k*_−,*e*_ = 4 *s*^−1^, *k*_ℎ*yd*_ = *k_Pir_* = 1.54 · 10^−3^*s*^−1^, *k*_+,*n*_ = 0.28 µ*M*^−1^. *s*^−1^ and *k*_−,*n*_ = 0.0141 *s*^−1^. Note that the model considers identical dynamics at both filament-ends and that intrafilament nucleotide exchange dynamics are independent of the nucleotide state. (E) HS-AFM time-course of untagged MreB (1.2 µM) filaments polymerized on an SLB (DOPC:CL 80:20) upon addition of 0.2 mM of ADP. Time in minutes:seconds. Scale bar, 200 nm. (F) Quantification of the effect of the ADP flush on preformed filaments. Standard deviation bars on top only. Data from two independent experiments. (G) Representative image of the capping experiment, in which E136A pFAST-MreB visualized by ATP* (green) was added to an existing filament of WT pFAST-MreB at t0. Both WT and E136E are labeled by TFCoral (red). Time in minutes:seconds. Scale bar, 1 µm. (H) Representative time-lapses of E136A-capped WT filaments labelled by ATP* when the buffer was replaced with MreB-free ADP-containing (0.2 mM) buffer. There are several seconds between the addition of ADP and the start of the movie. Blue arrows indicate breakage at the WT segments. Yellow arrows indicate the direction of depolymerization. (I) Simulation of the time evolution of MreB filament length, starting from a MreB filament of initial length of 5000 subunits and randomly bound to ATP, ADP-Pi, ADP, or nucleotide-free (bound probabilities *p_ATP_* = *p_ADP_*_−*Pi*_ = *p_ADP_* = *p_e_* = 0.25), in the absence of monomers. Green curve simulates the case where intrafilament nucleotide exchange does not occur (*k*_+,*n*_ = *k*_−,*n*_ = 0); Magenta curve simulates the case where intrafilament nucleotide exchange with free ADP (100 µM) occurs; Blue curve simulates the case where intrafilament nucleotide exchange with free ATP (100 µM) occurs. (J) CCCP-induced delocalization of MreB in growing cells visualized by TIRF microscopy. Cells expressing GFP-MreB WT (strain RCL421) and E136A (strain RCL2028) were exposed to 100 µM CCCP just before T0. Numbers above images correspond to time post T0, in minutes. Scale bar, 5 µm.

To explain the polymerization and disassembly dynamics of MreB filaments, we developed a stochastic Monte-Carlo model incorporating experimentally determined kinetic parameters for polymerization, depolymerization, ATP hydrolysis, Pi release and nucleotide exchange (Fig. 4D and Supplementary Information, ‘Modeling MreB dynamics’). The model accurately reproduces MreB polymerization kinetics in the presence of ATP (Fig. S8). A key difference between WT and ATPase mutant filaments is their nucleotide composition during polymerization: E136A filaments are only ATP-bound due to the absence of hydrolysis or nucleotide release; WT filaments remain mostly ATP-bound because of intrafilaments nucleotide exchange and relatively high ATP affinity; D158A filaments, however, are largely nucleotide-free because of their low affinity for ATP (Fig. S9A). Simulating a buffer exchange to MreB-free and nucleotide-free buffer (Fig. S10) mirrors the distinct behaviors observed experimentally (Fig. 4C). WT filaments disassemble irregularly, initially at *k*_−,*t*_ while filaments are ATP-bound, then accelerating as ATP is hydrolyzed to ADP or dissociates from subunits (Fig. S10A, B). E136A filaments remain stable, as ATP is not hydrolyzed and stays bound (Fig. S10D, E). Finally, D158A filaments, rapidly deprived of their bound nucleotides, disassemble immediately at a rate *k*_−,*e*_ higher than *k*_−,*t*_ (Fig. S10G, H).

We next sought to examine the function of intrafilament nucleotide exchange by replacing the solution after polymerization with MreB-free buffer containing ADP. WT MreB filaments underwent very rapid shrinkage (−70.34 ± 29.89 nm.s^-1^, > 5-fold faster than the maximum elongation rate), with all filaments disappearing from the TIRF field in less than 1 minute (Fig 4A,B and Movie 6). Similar ADP-induced disassembly was observed at higher filament densities in HS-AFM experiments (Fig. 4E and Movie 7). However, D158A filaments disassembled only marginally faster than in the presence of buffer alone, yet still at a much slower rate than WT MreB filaments (−22.79 ± 12.02 nm.s^-1^). E136A filaments remained stable and even exhibited slight positive elongation (1.80 ± 2.27 nm.s^-1^) (Fig. 4A,B,F and Movie 8), which could be due to the presence of few remaining ATP-bound MreB subunits on the SLB after the flush. In our simulations, presence of free ADP recapitulates again the filament behavior observed experimentally: E136A filaments remain stable, as ATP is not hydrolyzed and stays bound (Fig. S10D,F). D158A filaments, which bind nucleotides with low affinity, load poorly with ADP and depolymerize at a similar rate than in the absence of free nucleotide (Fig. S10G, I). WT filaments, on the contrary, load ADP efficiently. Once loaded with ADP, they depolymerize at a faster *k*_−,*d*_ rate (Fig. S10A, C).

Interestingly, free ADP was reported to also increase the rate of ParM filament disassembly, and this effect was proposed to result from nucleotide exchange at filament ends ^29^. To test whether MreB filaments in the presence of ATP become stabilized by an ATP cap or by intrafilament nucleotide exchange, we capped WT filaments labelled with TFCoral with E136A MreB labelled with both ATP* and TFCoral (t_0_ in Fig. 4G and Movie 9). When the solution was flushed with free ADP (and TFCoral to ensure continuous labelling of the filaments), filaments no longer underwent rapid (< 1 min) end-wise depolymerization. Instead, after less than 1 min, most filaments started breaking in the WT section or near one cap and depolymerized, leaving only the E136A caps (Fig. 4F-H). In a few instances, filaments remained stable or completely depolymerized (Fig. 4F, Movie 9), possibly because of statistical fluctuations in E136A cap sizes. These results show that ADP-rich MreB filaments disassembly occurs through both rapid depolymerization at rates corresponding to *k*_−,*d*_ and filament instability leading to fragmentation.

Altogether, our findings that ATPase mutants polymerize and that WT filaments exhibit slow hydrolysis support a model in which ATP hydrolysis is mechanistically uncoupled from polymerization. Instead, hydrolysis occurs after polymerization, and filament ends are capped by one or several ATP or ADP-Pi-bound subunits that stabilize the filament during polymerization (Fig. S9A). Furthermore, our model predicts that, because the rate of intrafilament nucleotide exchange is significantly faster than the rate of Pi release, the nucleotide state of MreB filaments (and thus their stability) is highly dependent on the pool of free nucleotides. To illustrate this significant result, we simulated the fate of WT MreB filaments randomly bound to ATP, ADP-Pi, or ADP, either in the absence of free nucleotide or in the presence of a free pool of ADP or ATP. In the presence of ADP, filaments aged and disassembled rapidly, whereas ATP stabilized them, effectively acting as a filament rejuvenation mechanism (Fig. 4I).

### Nucleotide-dependent filaments stability *in vivo*

Given the high intracellular concentration of MreB (well above the Cc) and the absence of identified canonical ABP-like proteins, nucleotide-dependent conformational changes appear to be as a major regulator of MreB filament polymerization and stability in the membrane. ATP binding acts as a switch to promote polymerization and ATP hydrolysis enables depolymerization, whereas intrafilament nucleotide exchange tunes filament stability on the membrane: exchange with free ATP in solution stabilizes the filaments, while exchange with free ADP promotes their disassembly. To test this *in vivo*, we monitored the localization of both MreB WT and E136A in exponentially growing *B. subtilis* cells upon addition of carbonyl cyanide m-chlorophenyl hydrazone (CCCP), which uncouples proton motive force and depletes the intracellular ATP pool. Before CCCP addition, cells expressing E136A at native levels displayed normal rod-shape, consistent with normal MreB function, and E136A localized in diffraction-limited motile patches similar to those observed for WT MreB (Fig. 4J and Fig. S11) ^7,29,32,33^. Upon CCCP addition, WT MreB filaments became progressively delocalized ^39^, while E136A filaments remained stable on the membrane, as predicted (Fig. 4J and Fig. S11). We propose that, in growing cells, high ATP levels maintain MreB filaments on the membrane through intrafilament nucleotide exchange, explaining their long lifespan ^5,12^. Under this model, normal ATPase activity is not required for MreB morphogenetic function. However, ATPase activity may be required for dynamic MreB relocalization during the cell-cycle, developmental processes or stress conditions that lower ATP levels. The similar affinities of MreB for both ATP and ADP then further suggest that the intracellular ATP/ADP ratio may ultimately control MreB function ^17^. Conditions that shift the ATP/ADP ratio shifts toward higher ADP levels include the transition to stationary phase, sporulation initiation, reduction of ATP synthesis (e.g. inhibition of respiration or electron transport chain), and environmental stresses that typically correlate with growth reduction, which in turn correlates with MreB-driven cell wall elongation.

In summary, the membrane-coupled, ATP-dependent polymer phase cycle of MreB defines a new class of self-organizing biological filament dynamics, highlighting an evolutionary solution for maintaining stable, membrane-bound filaments during growth.

## Materials and Methods

### Plasmids and strains construction

Primers and strains used in this study are listed in Supplementary Table S1 and Table S2, respectively. Full-length mreB from Bacillus subtilis was cloned in a pET28a vector (MilliporeSigma, Novagen) as translational fusion with a N-terminal 6xHistidine (6xHIS) tag for affinity purification. We also inserted a promiscuous Fluorescence-Activating and absorption-Shifting Tag (pFAST, gift from Arnaud Gautier) at the N-terminus 1. The sequence of the plasmid is provided in Supplementary Table S3. This pET28a 6xHIS-pFAST-MreB plasmid was used as a template to generate the two ATPase mutants E136A and D158A by site-directed mutagenesis using primer pairs 272/273 (E136A) and ac-1116/1117 (D158A). To produce untagged MreB, a pET28a-SUMO vector was ordered from Genecust (Boynes, France) containing the complete mreB open reading frame from B. subtilis codon-optimized for E. coli expression. pET28a-HsSENP1 encoding human SENP1 was ordered from Addgene (plasmid #71465).

To generate the B. subtilis strain expressing gfp-mreBE136A at the native mreB locus (strain RCL2028), primers pair CC272/273 was used to introduce the mutation, and primers CC181/RK14 were used as external primer, using as template genomic DNA from strain RCL0421, expressing a translational gfp-mreB fusion previously shown to be functional 2 and optimized to express wild-type levels of MreB. To generate the B. subtilis strain expressing pFAST-mreB at the native locus (strain RCL0925), the pFAST sequence was amplified from the pFAST plasmid (Supplementary Table S3) using primers pair CC592/CC564. The regions upstream and downstream mreB were amplified by PCR from strain RCL0421, using primers CC181/CC593 and primers CC565/CC182. After Gibson assembly, the product was amplified with external primers pair CC181/CC182.

### Protein purification

For protein expression and purification from the heterologous host E. coli, T7 express (New England BioLabs, N-E, USA) cells were transformed with the MreB constructs or SENP1. Bacteria were grown in 1 L of lysogeny broth (LB) supplemented with 25 µg/µl of kanamycin to an OD600 of 0.6 at 37°C, after which protein expression was induced with 1 mM Isopropyl β-d-1-thiogalactopyranoside (IPTG) over-night at 15°C, except for the expression of SENP1, which was induced with 0.1 mM IPTG for 3 h at 30°C. Bacteria were harvested by centrifugation (5000 g for 10 min at 4 °C). The pellet was washed once in buffer A (20 mM Tris-HCl pH7, 500 mM KCl) and stored at −70°C for later use. On the day of the protein purification, the pellet was thawed and resuspended in xx ml of buffer A supplemented with 0.25 mg/ml lysozyme and a half tablet of EDTA-free Complete protease inhibitors (Roche). For the purification of SENP1 the pellet was resuspended in 20 mM HEPES pH7.5, 500 mM KCl, 5 mM Imidazole, 10% glycerol, 0.1% triton, 2 mM β-mercaptoethanol and EDTA-free Complete protease inhibitors (Roche). The suspension was sonicated on ice with a Vibra-Cell VC505 processor (Sonics & Materials, Inc, Newton, CT, USA) for 12 min with 10/15 s on/off cycles at 50% power, and the supernatant was collected after clarification by centrifugation (40 000 g for 20 min at 4 °C).

The 6xHistidine pFAST-MreB constructs were purified following a 2-step procedure as previously described 3. Briefly, first the proteins were purified by affinity chromatography on a Ni-nitrilotriacetic acid (Ni-NTA) agarose resin (Thermo Fisher Scientific). Secondly, the proteins were loaded on a size exclusion chromatography HiLoad 16/60 Superdex 200 pg column (GE Healthcare Life Sciences / Cytiva), connected to an AKTA FPLC system (GE Healthcare Life Sciences). MreB-containing fractions were pooled and concentrated with an ultrafiltration spin column (Vivaspin, 10 000 MWCO; Sartorius). Concentration was determined from the absorption at 280 nm measured using a Nanodrop spectrophotometer (Thermo Fisher Scientific). The recombinant proteins were aliquoted and immediately frozen and stored at –70 °C.

SENP1 protein was purified using affinity chromatography on a Ni-NTA agarose resin (Thermo Fisher Scientific) similarly to MreB. SENP1 proteins were aliquoted and immediately frozen and stored at –70 °C.

For the production of tag-less MreB, after the first Ni-NTA affinity purification the 6xHIS-SUMO-MreB protein was incubated with purified SENP1 and put in over-night dialysis in buffer A supplemented with 1 mM DTT and 2 mM EDTA at 4 °C with gentile stirring to cleave the SUMO tag and remove the imidazole in a single step. The cleaved protein was pulled through a Ni-NTA resin to remove the His-tagged SENP1 and the cleaved-off His-SUMO tag. Cleaved MreB was further purified using size exclusion chromatography and processed similar to pFAST-MreB.

To determine whether recombinant MreB co-purified with a bound nucleotide, we measured the A260/A280 absorbance ratio and obtained a value of 0.62, indicating little or no ATP present. As a control, the addition of ATP at a 1:1 molar ratio with MreB increased the A260/A280 ratio to 1.26.

### Extraction of polar membrane lipids from B. subtilis

B. subtilis cultures were grown at 30°C in LB until OD600 0.5. Bacteria were harvested by centrifugation (5000 g for 10 min at 4 °C). The pellets were washed once with 0.9% NaCl, 0.02% Triton X100 and twice with 0.9% NaCl. The pellets were frozen and lyophilized overnight. The dried weight was carefully measured. The extraction of the polar membrane lipids was performed as described with modifications 4–6. The lyophilized pellets were treated with a chloroform, Methanol, 0.3% NaCl mixture with the ratio of 1:2:0.8 at 80°C for 15 min in glass tubes with Teflon caps. The extracts were vortexed for 1 h and centrifuged for 15 min at 4000 rpm at room temperature. The supernatant was collected and treated again with the Chloroform, Methanol, 0.3% NaCl mixture with the ratio of 1:2:0.8 at 80°C for 15 min. The extracts were vortexed for 30 min and centrifuged for 15 min at 4000 rpm at room temperature. The supernatant was collected and additional chloroform and 0.3% NaCl was added and the mixture was centrifugated at 4000 rpm for 15 min to achieve phase separation. The upper phase was discarded and chloroform phase was collected and transferred to a smaller glass tub. Finally, the chloroform was evaporated under a nitrogen stream and the lipid pellet was resuspended in chloroform to a concentration of 25 mg/ml and stored at −20°C. The lipid concentration was estimated based on the initial dry weight of the bacteria and the lipid fraction of 5.2 % for the dry weight of B. subtilis 7.

### Preparation of lipid monolayers and transmission electron microscopy

The lipid monolayers and the transmission electron microscopy (TEM) were performed as previously described 3 with some alterations. In short, MreB protein was added to 20 mM Tris pH7, 100 mM KCl, 2 mM ATP, and 5 mM MgCl2 and deposited as a 20 µl droplet on a plastic support. B. subtilis extracted lipids dissolved in chloroform (1 mg/ml) were added carefully to the top of the drop and incubated for 1 h at 37°C. Afterwards, carbon-coated electron microscopy grids (CF300-Cu from Electron Microscopy Sciences or 1753-F from Ted Pella, Inc) were placed on top of the grids with the carbon-side facing down for two minutes. The grids were stained with 1% uranyl acetate and air-dried. TEM images were acquired on a charge-coupled device camera (AMT) on a Hitachi HT 7700 electron microscope operated at 80kV (Milexia, France)8. Semi-quantitative TEM was performed, on images acquired at 30 000 x magnification, as previously described 3. Briefly, random fields of view were chosen on a TEM grid, before zooming in and recording for the presence of high (> 10 polymers) or low (1-10 polymers) polymer density or the absence of polymers.

### Liposome preparation

Liposomes were prepared by adding lipids dissolved in chloroform and dehydrating the lipid mixture first under a stream of argon gas and secondly in a desiccator under vacuum at room temperature for 3 h. Several lipid compositions were used (listed in Table S4). Lipids were rehydrated in either 25 mM HEPES pH 7.5, 300 mM KCl buffer or in 20 mM Citrate pH 4.6, 300 mM KCl buffer, for either the liposome binding assay or for formation of a Supported Lipid Bilayer (SLB), respectively. Liposomes were formed by vortexing and incubating at 40°C, or at 55°C for liposomes containing B. subtilis extracted lipids. Lipid mixtures were passed 21 times through a lipid extruder with a 100 nm pore size (Avanti Polar lipids) on a heated plate at 70°C (internal temperature 40°C) to produce small uniform-sized liposomes. A final concentration of 0.4 mM liposomes was used in case of SLB formation and 1 mM of liposomes for the liposomes binding assay.

### Liposome binding assay

MreB protein was first centrifuged at 35 000 rpm (TLA1.1) at 4°C for 10 min to remove protein precipitations. Next, 2 µM MreB in 25 mM HEPES pH 7.5, 300 mM KCl, 1 mM MgCl2, 0.2 mM ATP was incubated with liposomes at a final concentration of 0.7 mM at 25°C for 15 min. The reaction was centrifuged at 100 000 g for 25 min at 25°C to separate the soluble fraction from the liposome-containing pellet. The supernatant was mixed with 5x standard SDS sample buffer and the pellet was resuspended in 1x standard SDS sample buffer, both samples were boiled for 5 min. Samples were stored at −20°C until further use. The resulting fractions were loaded equally for SDS-PAGE (12% acrylamide). The gel was stained with Coomassie blue and imaged. The intensity of the bands was measured in ImageJ and the supernatant/pellet protein ratio was determined.

### Formation of Supported Lipid bilayers

Slides and coverslips were cleaned in advance by 30 min of sonication cycles, first in 2 mM NaOH, then in 100% ethanol and finally in ultra-pure water (MilliQ), with extensive rinsing with MilliQ in between steps. Next, tglassware were cleaned by plasma treatment on a PDC-002-CE plasma cleaner (Harrick, XX) for 10 min at high RF level, before assembling them into imaging chambers. The chambers were made by adding 3 thin strokes of 2 layers of parafilm in between the cleaned slide and coverslip and placing them on a hot plate until the parafilm melted and fused the two together, creating two channels. 3 mM CaCl2 was added to the liposomes to facilitate liposomes fusion before injecting them onto the chambers. The imaging chambers were placed in a humid chamber at 40°C overnight. The next morning the SLBs were washed with 25 mM HEPES pH 7,5 and stored at 40°C to be used that day.

### Single filament TIRF microscopy

Imaging buffer was made with a final concentration of 25 mM HEPES pH 7.5, 1 mM MgCl2, 100 mM KCl, 70 mM DTT, 20 mM glucose, 1 mM DABCO, and 66 units/ml Glucose Oxidase (Aspergillus niger, Sigma G7141) and 868 units/ml Catalase (Bovine liver, Sigma C1345). Nucleotides were added at various concentrations ranging from 200 µM to 50 µM. For visualizing MreB-pFAST 5 µM of either TFLime (P-480541-250, The Twinkle Factory or TFCoral (516600-250, The Twinkle Factory) was added. For visualizing MreB without a tag, or as an alternative for visualizing pFAST-MreB, 3 µM ATP-ATTO-488 was added to the 50 µM of ATP or used pure. MreB protein was centrifuged at 35,000 RPM TLA1.1 at 4°C for 10 min to remove protein precipitations. MreB was added to the imaging buffer and added to the imaging chamber. For the wash out or nucleotide change experiments, the buffer in the imaging chamber was replaced with buffer with or without 0.2 mM ADP. TIRF microscopy was performed on an inverted microscope (Nikon Ti-E) with an Apo TIRF × 100 oil objective (Nikon, NA 1.49), with an iLas2 laser coupling system from Gataca Systems (150 mW, 488 nm and 50 mW, 561 nm). Images were collected with an electron-multiplying charge-coupled device camera (iXON3 DU-897, Andor) at maximum gain setting (300) via a 2.5× magnifier, yielding a final pixel size of 64 nm.

### Nanofabrication of HS-AFM cantilevers

BL-AC10DS-A2 HS-AFM cantilever (Olympus, Tokyo, Japan) was used as the scanning probe to film dynamic MreB protein polymerization. The cantilever has a spring constant (k) of 0.1 N/m and a resonance frequency (f) of 0.6 MHz in water (1.5 MHz in air). Electron beam deposition (EBD) was performed to fabricate a cantilever by growing a long and sharp tip (small apical radius), which is important to obtain high-resolution images. Cantilevers were first cleaned using UV/O3 treatment and then soaked in the piranha solution containing sulfuric acid and hydrogen peroxide. Next, EBD was conducted (30 kV accelerating voltage, 2 min irradiation) using a field emission scanning electron microscope (ELS-7500, Elionix Inc., Tokyo, Japan).

### HS-AFM nanoimaging

HS-AFM nanoimaging was performed using a laboratory-built HS-AFM microscope, as previously reported 9,10. A laser beam (670 nm wavelength) focused on an EBD-processed cantilever using a 20x objective lens CFI S Plan Fluor ELWD, Nikon, Tokyo, Japan). Dynamic cantilever deflection was detected by sensing the position of the laser beam reflected by the cantilever with a position-sensing-two-segmented photodiode. An optimal tip-sample loading force was formed by adjusting the free oscillation amplitude of the cantilever (A0) to 1.5–2.5 nm and the set point to 80–90% of the free amplitude. A glass stage glued to a stack of muscovite mica layers was mounted on the HS-AFM scanner. Flat mica surface served as a substrate for sample adsorption. To visualize dynamic real-time MreB polymerization, mica needs to be coated with a lipid layer (DOPC:cardiolipin 80:20). First, poly-L-lysine was added on a freshly cleaved mica sheet, washed after about 10 minutes, and then loaded with previously formed liposomes. After about 20 minutes, lipid-coated substrate was washed with buffer. The substrate was then scanned under a near-physiological buffer (25mM HEPES KOH, 100mM KCl, 1mM MgCl2, 0.2mM ATP), MreB protein added sequentially into buffer chamber during scanning to observe polymerization.

### HS-AFM image processing and data analysis

HS-AFM images were processed and analyzed using the ImageJ software (https://imagej.nih.gov/ij/). A first-order polynomial fit in both x-and y-directions was performed, followed by convolution with a Gaussian smoothing function to suppress noise and improve image quality. To observe MreB doublets, image files were converted to 8-bit grayscale and subjected to Fast Fourier Transform (FFT) bandpass filtering, with large structures filtered at 8 pixels and small structures filtered at 1 pixel. HS-AFM images were either kept as stack image (tiff) or exported as AVI format movies.

### ATPase activity assay

ATPase activity of MreB was assayed by measuring the release of free inorganic phosphate (Pi) in a colorimetric assay using malachite green 11,12, as previously described 3. The reaction was initiated by the addition of MreB to the reaction mixture, composed by the reaction buffer (25 mM HEPES pH 7.5, 100 mM KCl, 1 mM MgCl2, 1mM DTT) with appropriate supplements (e.g. 0.5 mM ATP, 0.5 mg/mL liposomes). The liposomes were made on the day of the experiment by mixing DOPC and Cardiolipin following an 80/20 ratio, followed by desiccation and resuspension in milliQ water to 10 mg/ml. After a 2 h incubation at 37°C, the reaction was ended by addition of 1 reaction volume of malachite revelation buffer (0.2% (w/v) ammonium molybdate, 0.7 M HCl, 0.03% (w/v) malachite green, 0.05% (v/v) Triton X-100). The quantity of Pi produced was determined by measuring the absorbance at 650 nm on a 96-well plate spectrophotometer (Synergy 2, Biotek).

### QCM-D measurements

DOPC and cardiolipin 18:1 in chloroform were purchased from Avanti Polar Lipids Inc. (Alabaster, AL). We prepared lipid mixtures of DOPC/CL at a molar ratio 80:20 in cryogenic vials for a final concentration of 0.5 mg/mL. These lipid mixtures were gently evaporated using a stream of argon to form a lipid cake. The lipid cake was dried overnight under vacuum. Immediately prior to QCM-D measurements, the lipids were resuspended in isopropanol by carefully vortexing for 10 sec. We used a QCM-D E4 (QSense AB, Biolin Scientific AB, Gothenburg, Sweden) was used to measure B. subtilis MreB binding to supported planar lipid bilayer as previously reported 3,13. QCM-D measurements record frequency and dissipation changes based on the piezoelectric properties of the crystal probe 14. Commercially available quartz crystals (QSense AB, Biolin Scientific AB, Gothenburg, Sweden) were equipped with a 50-nm-thick layer of silicon dioxide by chemical vapor deposition (GeSiM GmbH, Dresden, Germany). The quartz crystals were thoroughly cleaned in a 1:1:5 volumetric ratio of concentrated ammonium hydroxide (Sigma-Aldrich), 30% hydrogen peroxide (Sigma Aldrich), and ultrapure water (Merck) at 70 °C. Prior to the measurement, the quartz crystals were oxidized in a plasma cleaner (Harrick Plasma, Ithaca, NY) for 3 min at high RF and immediately placed into the QCM-D chambers and HEPES-NaCl buffer (10 mM HEPES (pH 5.5), 100 mM NaCl) is added at a flow rate of 100 µL/min using a peristaltic pump. The measurement is started and after baseline stabilization, we followed the protocol for solvent-assisted lipid bilayer formation 15 (Figure S4, ‘1.’). Briefly, after approximately 10 min in HEPES-NaCl buffer, isopropanol is added at a flow rate of 100 µL/min for 10 min until the measurement signals stabilize. The addition of isopropanol leads to a sharp frequency and dissipation change. After 10 min, 0.5 mg/mL of the lipid mixture in isopropanol are added at 100 µL/min for another 10 min. The addition decreases the frequency by approximately −7 Hz (Fig. S4, ‘2.’). After another 10 min, the solvent exchange is started by adding HEPES-NaCl buffer to the isopropanol-lipid mixture for 20 min (Figure S4, ‘3.’), which leads to supported lipid bilayer formation at the previously reported characteristic frequency changes for spontaneous lipid bilayer formation 16. On this supported lipid bilayer, we studied the adsorption of B. subtilis MreB at varying concentrations of 0.1 µM, 0.5 µM, 1 µM and 5 µM to the bilayer at 100 µL/min. During MreB adsorption and desorption, MreB was almost completely displaced from supported lipid bilayers which allowed us to perform successive MreB adsorption measurements on a single supported lipid bilayer. Each adsorption measurement is done in 4 replicates and each measurement was repeated at least twice.

### Fluorescence Anisotropy experiments

Nucleotide binding experiments were initiated by adding a fluorescent nucleotide analogue N6-(6-Amino)hexyl-ATP-ATTO-488, N6-(6-Amino)hexyl-ADP-ATTO-488, or N6-(6-Amino)hexyl-AMPPNP-ATTO-488 (referred here to as N*; Jena Biosciences), to MreB in buffer (20 mM Tris pH 7, 500 mM KCl, 1 mM DTT, 2 mM EDTA and 5 mM MgCl₂). Binding of N* to MreB was monitored by measuring fluorescence anisotropy, using excitation and emission wavelengths of 504 nm and 521 nm, respectively 17. All measurements were carried out using a Safas Xenius XC spectrofluorometer (Safas Monaco) operated with SP2000 software (version 7.8.13.0).

For binding affinity measurements, steady-state fluorescence anisotropy values were determined at constant concentration of N* (0.2 µM) while increasing the concentration of MreB. Values were recorded 5 minutes after the beginning of the experiment. MreBBs was shown to be nuclotide free at the beginning of these experiments. Data were fitted and plotted using Igor Pro (version 9.0.5.1) in order to estimate the dissociation constant (Kd) according to the following equation:

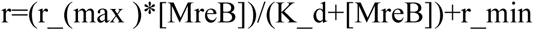

Where r corresponds to the measured fluorescence anisotropy, rmax represents the maximum anisotropy value when MreB is fully saturated with the fluorescent nucleotide (constrained between 0.2 and 0.23, the highest values usually measured for ATP* binding to MreB or G-actin 17, [MreB] is the concentration of the protein, Kd is the binding constant and rmin is the anisotropy value ofthe fluorescent nucleotide alone in solution (constrained between 0.035 and 0.045). Each data point represents the mean of at least three independent measurements, and error bars correspond to the standard deviations.

For kinetic analyses, the rate of fluorescent nucleotide binding at initial time points when MreB is mixed with N* is primarily governed by the rate of association of N* with MreB (konN*). In contrast, the reaction rate when N* is chased by excess ATP is controlled by the rate of dissociation of N* from MreB (koffN*). Monoexponential fits of these curves using Igor Pro (version 9.0.5.1) provided a first estimation of these 2 rate constants. We confirmed these values using Kintek Explorer version 1, using a simple model consisting of the two reactions:

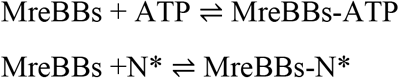

which involves four rate constants: konN*, konN, koffN and koffN*. Each reported rate constant represents the mean value obtained from at least three independent experiments.

### Sample preparation for in vivo microscopy (cell morphology and MreB dynamics)

B. subtilis strains were stored as glycerol stocks at −70°C. For each experiment, a small aliquot of thawed stock was used to inoculate an overnight preculture in LB at 30°C with shaking. Precultures of strains deleted for mreB, or expressing translational fusions GFP–mreB or pFAST–mreB, were supplemented with 20 mM MgSO4 and kanamycin (10 µg/mL); the wild-type preculture was grown in LB without supplements. Precultures were incubated overnight at 30°C with agitation. Overnight cultures were diluted into fresh LB to a low starting OD and grown at 37°C with shaking without antibiotics. Cultures were imaged during exponential phase. For “pFAST–mreB” samples, 40 µM TFLime dye was added to the culture, 30 min before imaging.

For imaging, 2% (w/v) agarose pads were prepared in LB, placed on a slide and prewarmed to 37°C. 4 µL of culture were deposited onto a prewarmed pad, covered with a glass coverslip to sandwich cells between pad and slide, and transferred immediately to the microscope stage to maintain temperature and cellular state 18.

For CCCP-induced delocalization experiments, cells expressing WT MreB (strain RCL421) and mutant E136A (strain RCL2028) were grown to mid-exponential phase of growth in rich LB medium at 37°C with strong aeration. A fraction of each culture was exposed to 100 µM CCCP (final concentration), marking the T0 of the experiment, and short 30 s timelapses in TIRF illumination (1s frequency, 488 nm laser, 100 ms exposure) were acquired every 15 to 20 min up to a maximum of 2h30. Control cells unexposed to the drug were acquired at T0. A maximum projection of each timelapse was generated and Images were displayed with a uniform intensity display range (min–max) set in Fiji for direct visual comparison; no intensity normalization was performed.

For live cell imaging to score the mobility of MreB patches, B. subtilis cells were first grown as overnight precultures at 30°C in LB with 20 mM MgSO4 and appropriate antibiotics, then diluted at an OD600 of 0.005 and grown at 37°C. Imaging was performed when cells reached early exponential phase (0.1 < OD600 < 0.2) and stationary phase (2.3 < OD600 < 2.5). One microliter of the liquid culture was then spotted onto an agarose pad (2 % in LB), topped by a coverslip and immersion oil, and mounted immediately in the temperature-controlled microscope stage set at 37°C. All acquisitions were done at 37°C, with an exposure time of 100 millisecond and interframe intervals of 0.5 s over 1 min. Time-lapse TIRFM movies were taken on a previously described setup 19.

### Wide-field epifluorescence and TIRF in vivo imaging (morphology and dynamics)

Imaging was performed on an inverted Nikon Ti E microscope equipped with an Apo TIRF 100× oil objective (NA 1.49). Excitation was provided by an iLas2 laser coupling system (Gataca Systems) with a 488 nm laser (150 mW). Fluorescence was recorded on an EMCCD camera (iXON3 DU 897, Andor) at maximum gain (300) via a 2.5× magnifier, yielding a final pixel size of 64 nm. Exposure time was 100 ms. Time lapse recordings for MreB dynamics were acquired for 1 min at 2 frames per second, using the Nikon Perfect Focus System to maintain focus. Image acquisition was controlled with MetaMorph v7.

### Image analysis

TEM images were taken randomly and either scored for the density of MreB protofilaments (none, low density or high density protofilaments, Figure 1A) or the length of MreB filaments protofilaments (Figure 1C). Filaments that were incompletely imaged were not measured, except if the filaments were spanning the entire length of the image.

SOAX software was used to analyze the curvature of MreB filaments of TIRF experiments 20,21. TSOAX was used to analyze the length of MreB filaments 21. TSOAX is an extension of SOAX (for network extraction in static images) to network extraction and tracking in time lapse movies. A script was made to extract the data with MATLAB to determine the rate of (de)polymerization by a linear fit to the filament-length versus time curve, using only the initial, linear region—where the fitting window was selected by visual inspection for each curve. Figures were made with PRISM software.

Fluorescence recovery after photobleaching (FRAP) experiment was performed on supported lipid bilayers made of DOPC + CL + TopFluor™ CL (79.8:20:0.2 molar ratios, table S1). Recovery curves were normalized to the pre-bleach and plateau fluorescence intensities, and the mobile fraction was determined from the extent of recovery. The lateral diffusion coefficient (D) was estimated from the half-time of fluorescence recovery (t₁/₂) according to the Soumpasis approximation, (0.224 · w2)/t1/2=D, where (w) is the radius of the bleached region. For a bleach radius of 2.3 μm and a recovery half-time of 3 s, D= 0.395 µm2/s. Mobile fraction = 0.7.

For cell diameter quantification, bright field images were segmented using Omnipose (version 1.0.6, custom model, 22). Cell width and length were measured as the dimensions of the minimum area rectangle enclosing each segmented mask.

For the histogram of the mean fluorescence intensity in Figure S3E, the mean fluorescent intensity was measured of 44 MreB filaments in FIJI (ImageJ) by manually placing a segmented line along the length of filaments and measuring the mean intensity.

### Statistical analysis

Statistical analysis was performed using PRISM software. Outliers were identified using PRISM software using the ROUT method with a False Discovery Rate (FDR), or Q, set to 0.2%. Statistical significance was assessed using a student’s t-test as indicated in the figure legends. Two-sided testing was used for each test. For student’s t-tests, data distribution was checked visually for normal distribution of the datapoints. For non-normal distribution statistical significance was assessed with Mann Whitney test.

## Acknowledgments

We thank former members of the ProCeD lab Aurélien Barbotin, Vlad Costache, Xavier Henry for their early contributions to this project, which ultimately evolved into the present study. We thank Martin Loose, Guillaume Romet-Lemonne, Juan Hermoso and Martin Alcorlo Pages, along all members of the ProCeD lab, for helpful discussions. We are grateful to Arnaud Gautier for the gift of pFAST tag and the fluorogenic ligands, to Alexandra Gruss for graciously lending her lab equipment and lipid extraction protocol, and to Adrià Sogues for the A_260_/A_280_ absorbance ratio check of our protein prep. We thank former ProCeD lab member Sarah Benlamara for some early technical assistance. We thank Christine Péchoux and Martine Letheule from the MIMA2 facility (Université Paris-Saclay, INRAE, AgroParisTech, GABI, 78350, Jouy-en-Josas, France) for TEM observations.

## Funding

European Research Council (ERC) under the Horizon2020 research and innovation program, ERC CoG No 772178 (R.C.L.)

Kanazawa University WPI-NanoLSI 2025 Bio-SPM Collaborative Research Program (IEA, RCL)

National Research Foundation (NRF) Singapore under its Mid-Sized Grant (MSG) NRF-MSG-2023-0001 (AM)

National University of Singapore through the Mechanobiology Institute A-0003467-00-00 (AM)

Microbes, DyMoB project 2025, Outgoing Mobility Grant, Center for Interdisciplinary Microbial Sciences, Paris-Saclay. (I.E.A., R.C.L.)

## Authors contributions

I.E.A: Conceptualization, Data Curation, Formal Analysis, Funding Acquisition, Investigation, Methodology, Visualization, Writing (original draft), Writing (review & editing).

C.B.: Data Curation, Formal Analysis, Investigation, Methodology, Resources, Writing (review & editing).

C.C.: Investigation, Writing (review & editing).

K.S.L.: Investigation, Writing (review & editing).

C.D.: Investigation, Writing (review & editing).

L.R.: Formal Analysis, Investigation, Methodology, Validation, Writing (review & editing).

C.P.C.: Investigation, Writing (review & editing).

A.J.: Methodology, Writing (review & editing).

R.W.W.: Investigation, Supervision, Validation, Writing (review & editing).

A.C.: Formal Analysis, Investigation, Writing (review & editing).

A.M.: Formal Analysis, Investigation, Methodology, Supervision, Validation, Writing (review & editing).

R.C.L.: Conceptualization, Formal Analysis, Funding acquisition, Project administration, Supervision, Validation, Writing (original draft), Writing (review & editing).

## Data, code, and materials availability

Research data will be deposited on Zenodo after publication and are available from the corresponding authors upon request. Constructs can be available from the corresponding authors under a material transfer agreement. Relevant identifiers are provided in the manuscript and Supplementary Materials.

## Supplementary Materials

Supplementary Text

MreB lipid membrane interactions QCM-D

Modeling MreB dynamics

Supplementary Figure Legends Supplementary Figures S1 to S11

Supplementary Tables S1 to S4 Movie captions

**Other Supplementary Material for this manuscript includes the following:**

Movies 1 to 9

