## Supplemental Materials for "Intrafilament nucleotide exchange in a prokaryotic actin homolog"

#### Supplementary Text

##### MreB lipid membrane interactions QCM-D

Previously, we generated supported lipid bilayers (SLBs) of various DOPC:DOPG mixtures by vesicle fusion on silicon surfaces [3]. Because this approach is limited to a narrow range of lipid compositions, we used solvent-assisted lipid bilayer formation [23] to prepare DOPC:cardiolipin (CL) SLBs. Briefly, the appropriate volume of lipid stock in chloroform (Avanti Polar Lipids) was suspended in a glass vial to reflect the desired molar ratio in chloroform. The total amount of lipid required for drying depends on the total volume of lipid sample solution needed for the experiment, in our case ~ 0.75 mg. The lipid solution was dried with a gentle stream of argon while slowly rotating the glass vial at a slightly tilted angle. The dried lipid films were left in a vacuum oven overnight at room temperature. Immediately before introducing the lipid sample into the measurement chamber, isopropanol was added to solubilize the dry lipid film, for a final lipid concentration of 0.5mg/mL. To ensure homogenization, the lipid solution was mixed by gently pipetting up and down for approximately 10 sec.

For quartz crystal microbalance measurements with dissipation monitoring (QCM-D), we used a QCM-D E4 (QSense AB, Biolin Scientific AB, Gothenburg, Sweden) as previously reported to measure lipid bilayer and protein interactions [3,24]. In QCM-D measurements, frequency and dissipation changes are recorded based on the piezoelectric properties and the adsorption kinetics on a crystal probe [14,25]. The quartz crystals (QSense AB, Biolin Scientific AB, Gothenburg, Sweden) were prepared with a custom 50 nm-thick layer of silicon dioxide by chemical vapor deposition (GeSiM GmbH, Dresden, Germany). The silicon dioxide crystals were thoroughly cleaned in a 1:1:5 volumetric ratio of concentrated ammonium hydroxide (Sigma-Aldrich), 30% hydrogen peroxide (Sigma Aldrich), and ultrapure water (Merck) at 70°C for 3 min prior to each measurement. The silicon dioxide crystals were immediately transferred to the measurement chambers of the QCM-D, which was set to 25°C. Peristaltic pumps were set up to inject HEPES–NaCl buffer (10 mM HEPES, 100 mM NaCl, pH 5.5) into the measurement chamber at a flow rate of 100 µL/min until the baseline was stable. After stabilization, isopropanol was pumped into the chambers at a fixed flow rate of 100µL/min for at least another 10 min to replace HEPES–NaCl buffer. After isopropanol completely replaced the HEPES–NaCl buffer, the 0.5mg/mL lipid mixture in isopropanol was introduced into the measurement chamber at a flow rate of 100 µL/min for at least 10 min until the measurement signals stabilized. This step resulted in frequency losses of ~4-7 Hz due to the adsorption of lipids on the silicon dioxide surface. In a final solvent replacement step, HEPES–NaCl buffer was pumped into the system at a flow rate of 100 µL/min for 20 min in order to enforce spontaneous SLB formation. This step ended in frequency (26-30 Hz) and dissipation ($0.1\text{-}1\times10^{6}$) changes that indicated successful SLB formation. After another 20 min, the SLBs were rinsed with MreB buffer (1 mM $MgCl_2$, 100 mM KCl, 10 mM HEPES pH7.5). At a stable baseline after solution exchange, MreB ($\pm$ 2 mM ATP, ADP, AMP-PNP) in was added at 0.1, 0.5, 1 and 2 µM (low to high concentration) to the SLB at a pump speed of 100 µL/min for 5 min. The adsorption of MreB was measured for at least 20 min before exchanging and rinsing with buffer for 5 min at 100 µL/min. In a series of experiments (from low to high MreB concentration), MreB was almost completely displaced by the rinsing step, allowing multiple adsorption steps on a single SLB. The analysis software QTools (QSense AB, Biolin Scientific AB, Gothenburg, Sweden) was used to calculate the thickness from frequency shifts using the Sauerbrey model. Each measurement was repeated at least twice with 4 repeats per run.

**Modeling MreB dynamics**

I. Objective of the model

The modeling work simulates the temporal polymerization/depolymerization dynamics of wild-type (WT), E136A and D158A mutant MreB filaments. The model accounts for the nucleotide-bound state of MreB filament subunits, the concentration of free MreB monomers and the concentration of soluble nucleotides (ATP or ADP). The goal is to identify a plausible biochemical model that reproduces experimental data, and to validate a set of biochemical rate constants that describes MreB dynamics.

II. Biochemical reactions at play

The model takes into account the 6 following reactions, which are shared with eukaryotic actins, albeit at different rates (Fig. 4D):

- one polymerization reaction, corresponding to the addition of ATP-bound MreB monomers to the ends of MreB filaments, of rate constant $k_{+,t}$. The addition of ADP-bound MreB monomers is not considered, as ADP-MreB monomers do not polymerize in vitro (Fig. 1C).
- three depolymerization reactions, which can occur when a terminal subunit of MreB filaments are bound to ATP, ADP-Pi or ADP. The corresponding rate constants are $k_{-,t}$, $k_{-,dp}$ and $k_{-,d}$.
- one ATP hydrolysis reaction, occurring stochastically on MreB subunits after polymerization, at a rate $k_{hyd}$. Inorganic phosphate is not released immediately after hydrolysis.
- one phosphate release reaction, occurring stochastically on MreB subunits after ATP hydrolysis, at a rate $k_{Pir}$.

The model also takes into account 3 MreB-specific reactions:

- one nucleotide dissociation rate along the filaments, occurring stochastically on MreB subunits, at a rate $k_{-,n}$.
- one nucleotide association rate along the filaments, occurring stochastically on MreB subunits, at a rate $k_{+,n}$.
- one depolymerization reaction, which can occur when a terminal subunit of MreB filaments is not bound to any nucleotide. The corresponding rate constant is $k_{-,e}$.

Importantly, because both ends of the filaments are identical, polymerization and depolymerization reactions occur at similar rates at both filament ends. The model does not take into account the double-stranded nature of MreB filaments, assuming that terminal subunits must polymerize or dissociate alternatively between strands to maintain filament stability. Thus, the model simulates filaments as a 1D chain of consecutive subunits.

III. Computational methods

Stochastic simulations use a Monte Carlo method, implemented with a rejection-free algorithm similar to the Gillespie algorithm [26,27]. In these simulations, all polymerization/ depolymerization, nucleotide hydrolysis, phosphate release and nucleotide exchange reactions are treated as probabilities for these events to occur.

The algorithm works by selecting the next polymerization/depolymerization event at each step, based on its probability of occurrence. The model runs as follows at each step:

1. The transition rates $r_i$ for each polymerization/depolymerization reaction at filament ends are calculated. For the association of MreB monomers, they are equal to $k_{+,t} \cdot [MreB - ATP]$. For the dissociation of MreB subunits, they are equal to $k_{-,t}$, $k_{-,dp}$, $k_{-,d}$ and $k_{-,e}$.
2. The sum of the transition rates $r_0 = \sum_i r_i$ is calculated.
3. One of the possible events is selected randomly, based on its probability $r_i/r_0$ to occur
4. The state of the filament is updated.
5. A uniform number u ∈ (0.1) is selected randomly and the time $t = t + \Delta t$ is updated (initial time is t = 0), where $\Delta t = -\log {}^{u}/_{r_0}$.
6. The simulation proceeds to random hydrolysis of ATP-bound MreB subunits, random phosphate release of ADP-Pi-bound MreB subunits and random nucleotide dissociation/association along MreB subunits based on the probability of these events occurring during a time interval $\Delta t$.
7. This process is repeated for a determined period of time or until the system reaches steady-state.

Simulations can provide various outputs such as the MreB filaments length or filament nucleotide composition over time. Simulations are performed with RStudio version 2026.01.0+392. Plots are generated with ggplot2.

#### IV. Initial set of rate constants deduced from experimental observations

While the model can explore a broad range of kinetic constant values, we initially constrained it using our experimental data, based on the assumptions outlined in this section.

##### *a. Association rate constant* $k_{+,t}$

The association rate constant $k_{+,t}$ is typically determined from the slope of the curve representing the elongation rate of filaments as a function of the MreB monomer concentration (Fig. 2E). At low monomer concentration (for $c < c_{sat} \sim 0.06 - 0.09$ µ$M$), elongation rates increase linearly with concentration, indicating a regime in which monomer diffusion to filament ends is rate limiting. In this regime, a linear fit of the experimental data provides a value of $k_{+,t} \approx 45$ µ$M^{-1}s^{-1}$, which is equal for WT, E136A and D158A mutant proteins. At high monomer concentration (for $c > c_{sat}$), elongation rates progressively saturate to a maximal value, indicating a regime in which monomer diffusion is not rate limiting anymore. Because MreB, unlike eukaryotic actin, has the property of polymerizing on a 2D lipid bilayers, one possibility is that MreB monomers must pre-bind to the membrane before polymerizing. Thus, a secondary membrane-associated and polymerization-competent MreB reservoir could have a lower diffusion rate or be depleted rapidly, reducing polymerization efficiency at a lower monomer concentration than observed for eukaryotic actins. This phenomenon is not included in the model, which is therefore only valid for MreB monomer concentrations below $c_{sat}$. Above $c_{sat}$, the elongation of MreB filaments occurs as if the apparent concentration were equal to $c_{sat}$.

We verified that the model recapitulates correctly the elongation of MreB filaments at different MreB monomer concentrations (Fig. S8A,B).

b. *Dissociation rate constant* $k_{-,t}$ *and critical concentration* $c_{c,ATP}$

The dissociation rate constant, $k_{-,t}$, and the critical concentration of assembly of MreB, $c_{c,ATP}$, can be deduced from the same curve representing the elongation rate of filaments as a function of the MreB monomer concentration. Intersection of the curve with the x axis (y = 0) indicates $c_{c,ATP} \approx 0.003\ \mu M$, while intersection with the y axis (x = 0) leads to $k_{-,t} \approx 0.135\ s^{-1}$. With this value of $k_{-,t}$, simulations predict accurately that pre-formed filament maintain the same average length for a MreB monomer concentration of 0.003 µM (Fig. S8C). $k_{-,t}$ and $c_{c,ATP}$ values are the same for WT, E136A and D158A proteins.

c. *Dissociation rate constant* $k_{-,dp}$

The disassembly rate of MreB filaments was not measured in this study in the presence of high inorganic phosphate concentrations that could saturate MreB binding sites. We rely here on observations from eukaryotic actin to estimate here that $k_{-,dp} \approx k_{-,t}$ (Table 1) for WT, E136A and D158A proteins.

d. *Dissociation rate constant* $k_{-,d}$

The dissociation rate constant $k_{-,d}$ can be estimated from the depolymerization rate of "aged" (ie. fully ADP-bound) MreB filaments. It is difficult to know the moment when MreB filaments are mainly bound to ADP in our experiments, but we can estimate that the fastest measured depolymerization rates (≈ 175 nm.s$^{-1}$ (Fig. 4B) in the presence of free ADP in solution) correspond to cases where MreB filaments are most saturated with ADP. Thus, considering 340 subunits per µm, we performed simulations for values of $k_{-,d} = 60\ s^{-1}$.

e. *Dissociation rate constant* $k_{-,e}$

Since nucleotide exchange occurs along MreB filaments, our model includes an intermediate state in which MreB subunits are not bound to any nucleotide. Thus, a dissociation rate constant $k_{-,e}$ is introduced to take into account the rate at which nucleotide free terminal subunits would dissociate from the ends of the filament.

Accounting for this intermediate state is particularly important for understanding the behavior of the D158A mutant, and is justified by the following observations: 1/ D158A mutant binds nucleotides with lower affinity than WT monomers (Fig. S7D,E) (although we acknowledge that a difference in affinity for monomers does not necessarily imply a difference in affinity for filament subunits); 2/ D158A filaments disassemble more rapidly than WT MreB filaments even in the absence of free nucleotide (Fig. 4B), raising the possibility that the nucleotide state of WT and D158A filaments is not identical after polymerization; 3/ D158A filaments polymerized in the presence of ATP-ATT0-488 are dimer than WT filaments, suggesting that a fraction of bound nucleotides dissociate from the filaments after polymerization.

The value of $k_{-,e}$ can be estimated from the depolymerization rate of the low nucleotide affinity mutant D158A. We considered that the depolymerization rates measured in absence of monomers and free nucleotides (Fig. 4B; ≈ 15 nm.s$^{-1}$) correspond to cases where D158A MreB filaments are least bound to a nucleotide. Considering 340 subunits per µm, we performed simulations for values of $k_{-,e} = 4\ s^{-1}$.

We can verify in a simulation that MreB filaments fully bound to ADP or nucleotide free, in the absence of ATP-bound MreB monomers, depolymerize completely at rates corresponding to $k_{-,d}$ and $k_{-,e}$, respectively (Fig. S8D). A 5000 subunit-long MreB-ADP filament depolymerizes in about 40 seconds, as observed experimentally for the fastest depolymerizing filaments (Fig. S8D).

*f. Nucleotide hydrolysis $k_{hyd}$ and phosphate release $k_{Pir}$ rates*

The experimental data provide a global quantification of the inorganic phosphate production rate during MreB polymerization ($k_{tot,Pir} = 0.046\ min^{-1} = 7.7 \cdot 10^{-4}\ s^{-1}$). Since this rate is calculated based on the total MreB concentration rather than the concentration of MreB that actually polymerized during the experiment, this value likely represents a lower bound of the true value. We explored here a range between $7.7 \cdot 10^{-2}\ s^{-1} > k_{tot,Pir} > 7.7 \cdot 10^{-4}\ s^{-1}$.

Moreover, $k_{tot,Pir}$ encompasses both the hydrolysis and the phosphate release kinetics from the filaments. Thus, each of these parameters cannot be determined independently, but only relative to one another. However, since the current model assumes identical polymerization/ depolymerization kinetics for ATP- and ADP-Pi-bound MreB subunits (see sections c., d. and f. of this Part IV.), ATP and ADP-Pi states remain equivalent until further development of the model. We can define a relationship between $k_{hyd}$ and $k_{Pir}$, as a function of $k_{tot,Pir}$. For two consecutive irreversible reactions:

$$MreB - ATP \xrightarrow{k_{hyd}} MreB - ADPPi \xrightarrow{k_{Pir}} MreB - ADP$$

the average time taken for MreB-ATP to transition to the MreB-ADP state equal:

$$\frac{1}{k_{tot,Pir}} = \frac{1}{k_{hyd}} + \frac{1}{k_{Pir}}$$

This equation defines how $k_{hyd}$ can be fixed as a function of $k_{Pir}$, and we ran our simulations here with $k_{hyd} = k_{Pir} \in [1.54 \cdot 10^{-1}, 1.54 \cdot 10^{-3}]\ s^{-1}$.

Contrary to WT, E136A and D158A are ATPase null mutants. Thus, $k_{hyd,E136A} = k_{hyd,D158A} = 0\ s^{-1}$. $k_{Pir,E136A}$ and $k_{Pir,D158A}$ values do not matter as no ATP is being hydrolyzed by the two mutants.

*g. Rate of intra-filament nucleotide exchange $k_{ex,F}$*

Based on our observations, we consider the possibility that intra-filament bound nucleotides dissociate from filaments and exchange with other nucleotides present in solution (Fig. 3F,H and I). The model assumes that MreB subunits can release their bound nucleotide regardless of its state (ATP, ADP-Pi or ADP), but can only bind a new ATP or ADP (not ADP-Pi). At the moment, we do not know if the rate of nucleotide exchange depends on the type of nucleotide exchange or not;

however, because of the similar affinity of ATP and ADP for MreB monomers, we assume a single intra-filament nucleotide exchange rate $k_{ex,F}$, which is different for WT MreB and its mutants but similar for ATP, ADP-Pi or ADP. $k_{ex,F}$ value is accessible experimentally from a monoexponential fit of our data (Fig. 3F).

$k_{ex,F}$ is a single parameter that reflects the kinetics of a two-step mechanism (dissociation of the previously bound nucleotide, followed by the binding of the subsequent one) with:

$$k_{ex,F} = \frac{k_{-,n}\, k_{+,n}[N]}{k_{-,n} + k_{+,n}[N]}$$

Where [N] is the concentration of free nucleotides. $K_{d,n} = k_{-,n}/k_{+,n}$ is the dissociation constants of the nucleotides from the side of the filaments.

Importantly, E136A mutant filaments do not exchange nucleotides (Fig. 3I). Thus, $k_{ex,F,E136A} = k_{-,n,E136A} = 0\ s^{-1}$ (the value of $k_{+,n,E136A}$ is not important for E136A because filament subunits are always bound to a nucleotide).

For WT filaments, our experiments measure $k_{ex,F,WT} = 0.014\ s^{-1}$ for [ATP] = 100 µM. Although $k_{-,n,WT}$ and $k_{+,n,WT}$ cannot be determined individually from our experiments, the relatively high affinity of WT MreB subunits for their nucleotide suggests that the rate intra-filament nucleotide exchange is mainly controlled by the dissociation constant $k_{-,n,WT}$. Hence, our simulations are initially based on a value of $k_{-,n,WT} = 0.0141\ s^{-1}$ (rate limiting) and $k_{+,n,WT} = 0.028\ \mu M^{-1} \cdot s^{-1}$ (fast association). With these values, $k_{ex,F,WT} = 0.014\ s^{-1}$ and $K_{d,WT} = 0.5\ \mu M$.

For D158A filaments, we do not have any experimental evidence whether intra-filament nucleotide exchange is controlled by the association constant $k_{+,n,D158A}$ or $k_{-,n,D158A}$. The values presented in this work are $k_{-,n,D158A} = 0.2\ s^{-1}$, $k_{+,n,D158A} = 10^{-4}\ \mu M^{-1} \cdot s^{-1}$, for which $k_{ex,F,D158A} = 0.01\ s^{-1}$ and $K_{d,D158A} = 2\ mM$.

Ultimately, these values reflect the experimentally observed differences in affinity, with $K_{d,E136A} < K_{d,WT} < K_{d,D158A}$ .

#### V. Experiments modeling to understand how nucleotides regulate MreB dynamics

This section analyzes the polymerization and flush experiments, in which MreB filaments, are initially polymerized in the presence of ATP (100 µM) and ATP-bound monomers, are later flushed into a new buffer. This buffer may or may not contain MreB monomers, and may or may not contain ADP (200 µM).

##### *a. Nucleotide state of WT and mutant MreB filaments during polymerization*

The first part of the experiment consists in the polymerization of the filaments in the presence of MreB monomers and ATP (100 µM). We simulated the polymerization of WT and mutants MreB filaments up to a size of 5000 subunits (ie. about 15 µm long filaments), using the full set of rate constants defined in IV (Fig. S9).

We analyzed first the nucleotide state of the polymerizing filaments in these conditions. Despite constant hydrolysis and Pi release, WT MreB filaments were found highly charged in ATP (> X%), the remaining subunits being mostly ADP-Pi (Fig. S9A). We assumed that, with a rate of intra-filament nucleotide exchange $k_{ex,F,WT} = 0.014\ s^{-1}$ higher than the rate of $k_{tot,Pir} = 7.7 \cdot 10^{-4}\ s^{-1}$, the recharging of ADP-bound MreB subunits after hydrolysis and Pi release was efficient. Indeed, decreasing $k_{ex,F,WT}$ by a factor 100 (with $k_{-,n} = 1.41\ 10^{-4}\ s^{-1}$) (Fig. S9B), or increasing $k_{tot,Pir}$ by the same factor (with $k_{hyd} = k_{Pir} = 1.54 \cdot 10^{-1}\ s^{-1}$) (Fig. S9C) decreases the proportion of bound ATP in the filaments. E136A filaments, which do not hydrolyze ATP nor exchange nucleotides, were found completely bound to ATP (Fig. S9A). D158A filaments, which do not hydrolyze ATP but which are weakly bound to their nucleotides, were found only partially bound to ATP, the remaining subunits being principally deprived of nucleotide (Fig. S9A).

Despite their different nucleotide composition, WT and MreB mutants were found to elongate at similar rates (Fig. S9D). For WT and E136A, which are mainly bound to ATP, this was expected. However, this was more surprising for D158A filaments which are largely deprived of ATP. Nevertheless, when analyzing the nucleotide state of terminal subunits of polymerizing D158A filaments (Fig. S9A), we noticed that those remain largely bound to ATP, thereby protecting these ends of the filaments from depolymerization. We compared the rates of subunit association to the filaments ($k_{+,t}\ [MreB] = 45 \cdot 0.05 = 2.25\ s^{-1}$) to the rate of nucleotide dissociation from the filaments ($k_{-,n,D158A} = 0.0141\ s^{-1}$) and found that, under these conditions, MreB monomers associate on average faster than nucleotides dissociate.

##### *b. Flush to a nucleotide-free buffer: case of WT MreB*

We analyzed then the depolymerization experiments, in which MreB filaments are flushed into buffers deprived of MreB monomers. In all these simulations, the simulated filaments are at the onset of depolymerization in the same nucleotide state as they were after their polymerization (see V. a).

We first sought to determine why WT MreB filaments, after a buffer flush removing ATP-bound MreB monomers and free nucleotides, depolymerize initially slowly (at about 1 $nm.s^{-1}$, ie. 0.34 $s^{-1}$ if we consider that MreB filaments contain 340 subunits.$\mu m^{-1}$; Fig. 4B), and then disappear from the membrane within 20 min. We simulated the depolymerization of the 5000 subunit MreB filaments in absence of free MreB monomers and free nucleotides (Fig. S10A, green). Contrary to ADP-bound MreB filaments (Fig. S8D), ATP-bound MreB filaments initially depolymerized slowly for about 250 s. After 250 s, they progressively depolymerized more rapidly until full depolymerization after about 1000 s. To analyze the mechanism underlying this acceleration, we studied the evolution of the filaments' nucleotide state over time (Fig. S10B). Initially, MreB subunits were primarily bound to ATP, and depolymerized at a rate of 0.28 $subunits.s^{-1}$ which corresponds to $2 * k_{-,t}$ (because filaments depolymerize symmetrically at both ends). Over time, MreB subunits mainly released their bound nucleotide, rather than hydrolyzing ATP it into ADP. Thus, the acceleration of depolymerization in these conditions is mainly driven by the rapid loss of the bound nucleotide at $k_{-,n,WT} = 0.0141\ s^{-1}$, rather than by the slower nucleotide hydrolysis at

$k_{tot,Pir} = 7.7 \cdot 10^{-4}\ s^{-1}$, and the depolymerization rate of MreB filaments evolves over time as a fraction of their subunits becomes nucleotide-free.

##### c. *Flush to a nucleotide-free buffer: case of E136A and D158A mutants*

The case of the E136A mutant is trivial. In absence of nucleotide release in solution nor ATP hydrolysis, filaments initially loaded with ATP remain bound to ATP and stable over time (Fig. S10D, green and F).

We studied then the case of D158A. We found that these filaments, initially largely devoid of bound nucleotides, progressively lose the remaining nucleotides (Fig. S10H). Hence, these filaments rapidly depolymerize at a rate corresponding to $k_{-,e}$ (Fig. S10G, green), as observed experimentally (Fig. 4B).

##### d. *Flush to an ADP buffer: case of WT MreB*

We then sought to determine the molecular mechanism which triggers the fast disassembly of MreB-WT filaments in the presence of ADP in solution. We simulated the depolymerization of ATP-bound MreB filaments in the absence of free MreB monomers but in the presence of free ADP in solution (Fig. S10A, magenta). We observed that filaments progressively switch from their ATP state into an ADP state (Fig. S10C). As a consequence, the depolymerization of filaments occurs notably faster, with an initial slow phase of about 200 s, and a total depolymerization time of about 400 s. This result reflects that intra-filament nucleotide exchange accelerates efficiently filament ageing, ie. the proportion of ADP-bound MreB subunits.

Nevertheless, these simulations do not fully capture the rapid, minute-scale depolymerization of the MreB filaments observed experimentally, due to the initial slow phase corresponding to ATP unloading from the filaments. One possibility is that ADP-rich filaments may also gradually become unstable and susceptible to fragmentation, which would accelerate their depolymerization. This phenomenon is not taken into account in these simulations but is clear in the E136A capping experiments (Fig. 4G).

##### e. *Flush to an ADP buffer: case of E136A and D158A mutants*

Logically, E136A MreB filaments are also stable in the presence of ADP due to the absence of nucleotide exchange for this mutant $k_{Ex,F} = 0\ s^{-1}$. Simulations indicate a slow and constant depolymerization of MreB filaments (Fig. S10D, magenta), at a rate corresponding to $k_{-,t}$, regardless of the presence of ADP or not. The slight polymerization observed experimentally in these conditions likely reflects the remaining presence of non-flushed ATP-bound MreB monomers on the membrane.

We next sought to understand why D158A filaments depolymerize only marginally faster in the presence of ADP than in its absence (Fig. 4B). Simulations indicate that the low affinity of these filaments for nucleotides only weakly promotes ADP binding (Fig. S10I). As a result, filaments remain mostly nucleotide-free and disassemble only marginally faster in our simulations too (Fig. S10I).

VI. Function of intra-filament nucleotide exchange

*a. Rate of ATP hydrolysis and Pi release versus rate of intra-filament nucleotide exchange*

Notably, the model reveals that the depolymerization rate of MreB filaments evolves over time as the fraction of each bound nucleotide $(f_t, f_{dp}, f_d, f_e)$ changes, with an overall kinetic constant $k_{-} = (f_t + f_{dp})\, k_{-,t} + f_d\, k_{-,d} + f_e\, k_{-,e}$. The fraction of bound nucleotide is controlled by two mechanisms which are ATP hydrolysis and Pi release, and intra-filament nucleotide exchange. Because these two mechanisms occur with different kinetics and different efficiency on WT and mutant MreBs, the nucleotide composition of WT and mutant MreB filaments is changing differently over time, thereby leading to different depolymerization dynamics.

A key finding of this work is the faster rate of intra-filament nucleotide exchange, at the nucleotide concentration used in these assays, compared to the rate of hydrolysis and phosphate release ($k_{ex,F,WT} = 0.0141\ s^{-1}\ vs\ k_{Pir} = 1.54 \cdot 10^{-3}\ s^{-1}$), which indicates that intra-filament nucleotide exchange promotes filament ageing faster than hydrolysis and nucleotide release. This result was suspected from our experimental measurments (Fig. 3A, F, H and I), and is confirmed by the fact that MreB filaments depolymerize at very different rates in the absence or presence of free ADP in solution. In the absence of ADP, the fact that filaments depolymerize very slowly indicates that their ADP-bound fraction remains low, while in the presence of ADP, the fact that filaments depolymerize fast indicates that their ADP-bound fraction is high.

*b. Intra-filament nucleotide exchange as a tuning mechanism of filament stability*

Because intra-filament nucleotide exchange is such an efficient mechanism, we questioned its function for controlling the ageing of MreB filaments. We simulated the depolymerization of a MreB filament initially set in a random nucleotide state, where each subunit has a probability $p_{ATP}$, $p_{ADP-Pi}$, $p_{ADP}$ or $p_e$ of being in a given nucleotide state (with $p_{ATP} + p_{ADP-PiP} + p_{ADP} + p_e = 1$). In the absence of intra-filament nucleotide exchange (corresponding to the case where filament ageing depends solely on nucleotide hydrolysis and Pi release), a 5000-subunit filament of initial composition $p_{ATP} = p_{ADP-Pi} = p_{ADP} = p_e = 0.25$ is predicted to disassemble in about 2500 s (Fig. 4I, green curve). In the presence of free ADP, intra-filament nucleotide exchange acts as a complementary filament ageing mechanism, and filaments disassemble in only 360 s (Fig. 4I, magenta curve). On the contrary, in the presence of free ATP, intra-filament nucleotide exchange acts as a rejuvenation mechanism, where continuous exchange for fresh ATP along MreB filaments stabilizes them and prevents their fast depolymerization (Fig. 4I, blue curve). Taken together, our data suggest that intra-filament nucleotide exchange serves as a mechanism complementary to nucleotide hydrolysis, regulating filament ageing based on the pool of nucleotide available in solution.

Figure S1

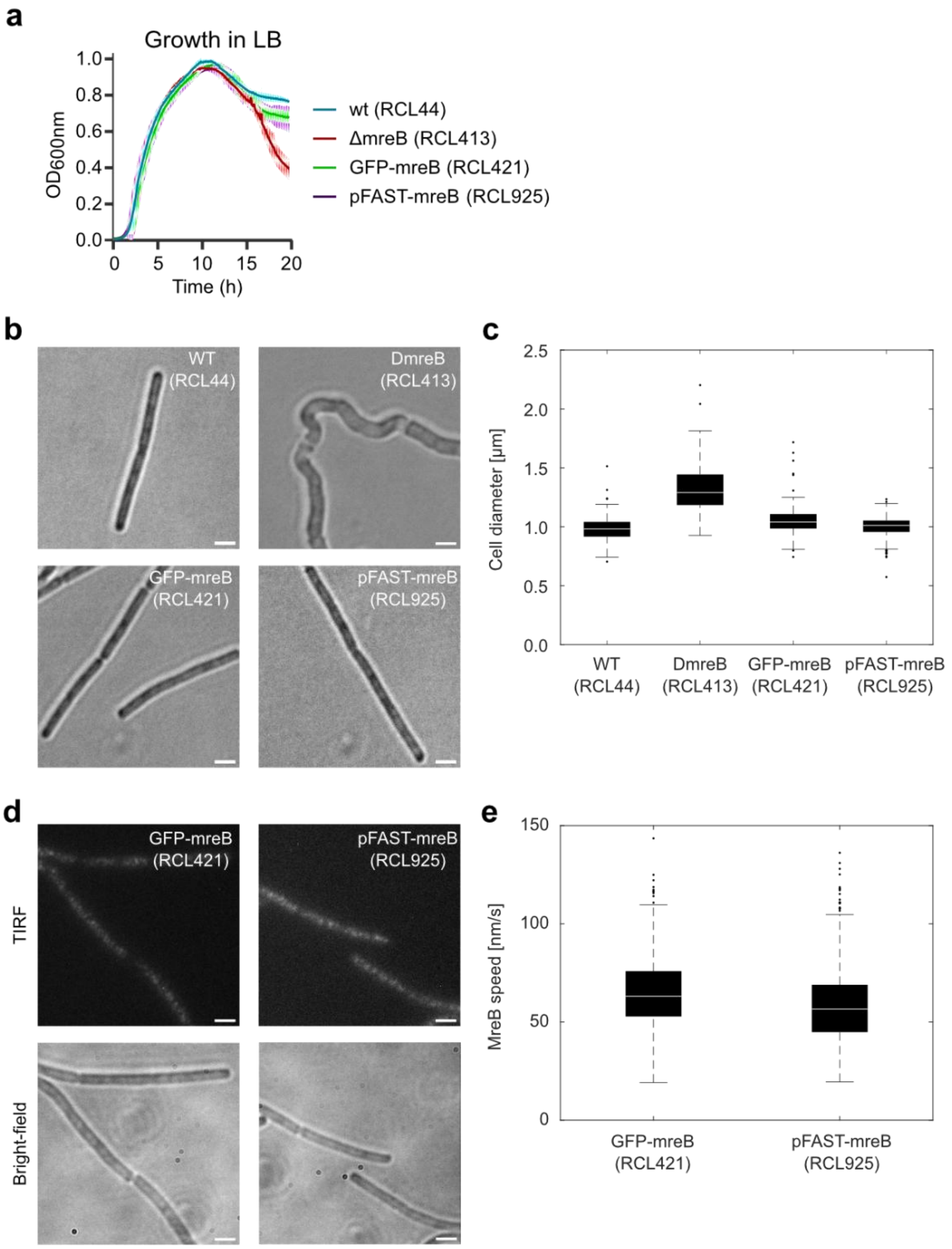

### Supplementary Figure and Legends

**Fig. S1. Functionality of the pFAST-MreB fusion *in vivo*.**
(A-C) Cells expressing *pFAST-mreB* at the native locus, as only copy *mreB* in their genome, display growth and morphology comparable to wild-type cells and to cells expressing a translational *gfp-mreB* fusion previously shown to be functional [19] and optimized to express wild-type levels of MreB [28]. Growth curves (A), representative wide-field epifluorescence images (B) and cell width distribution (C) of *B. subtilis* wild-type (strain RCL44), Δ*mreB* (RCL413), GFP-MreB (RCL421) and pFAST-MreB (RCL925) grown in LB. Scale bars, 2 µm. Widths (n > 100;

two independent experiments) were measured from wide-field images using the procedure detailed in Materials and Methods. Boxplots show the median, first and third quartiles; whiskers extend to the most extreme data points not considered outliers. Cells expressing GFP-MreB and pFAST-MreB display diameter and morphology comparable to the WT. WT, n=147; Δ*mreB*, n=103; GFP-MreB, n=166; pFAST-MreB, n=164. (D) Representative TIRF images showing the localisation of GFP–MreB and TFLime-labelled pFAST-MreB in growing cells, and corresponding brightfield epifluorescence images. Scale bars, 2 µm. (E) GFM-MreB and pFAST-MreB display similar localization and dynamics. Quantitative comparison of the speed of circumferentially moving MreB filaments measured by kymograph analysis. Boxplots show the median, first and third quartiles; whiskers extend to the most extreme data points not considered outliers (n > 1000; two independent experiments). Measured MreB speeds are comparable between the two strains. GFP-MreB, n=1251; pFAST-MreB, n=1061.

#### Figure S2

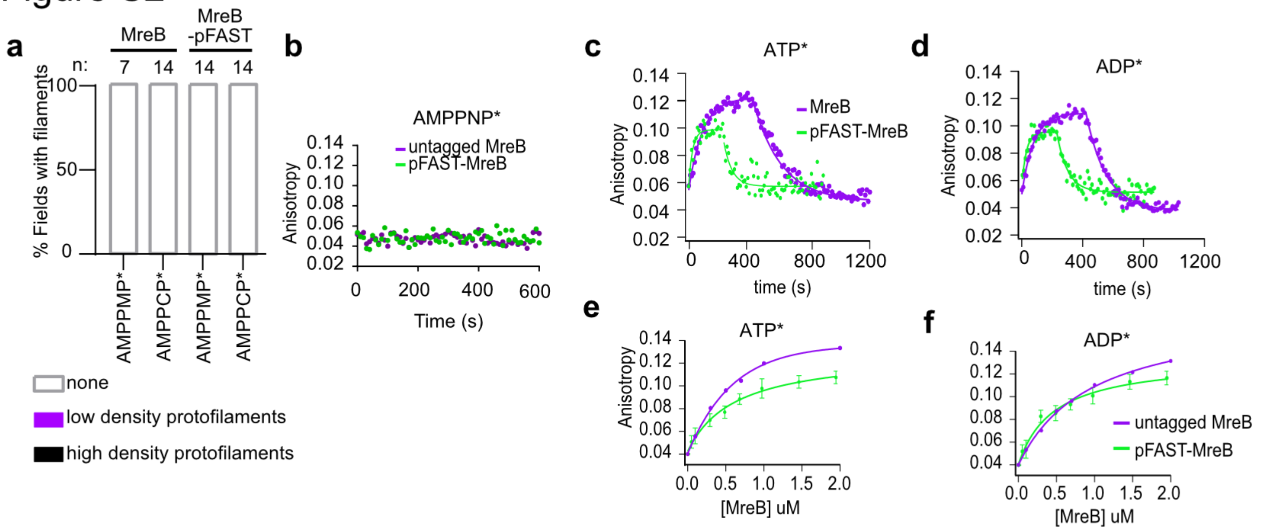

**Fig. S2. Analysis of filaments visualized by TEM and fluorescence anisotropy of MreB.**
(A) No filaments were detected on a lipid monolayer of *B. subtilis* lipid extract when MreB (1.5 µM) was incubated in the presence of 0.5 mM AMPPNP or 0.5 mM AMPPCP. n, amount of TEM fields analyzed. (B-D) Binding kinetics of 0.2 µM $N^6$(6-Aminohexyl)-AMPPNP-Atto-488 (AMPPNP*) (B), $N^6$(6-Aminohexyl)-ATP-Atto-488 (ATP*) (C) and $N^6$(6-Aminohexyl)-ADP-Atto-488 (ADP*) (D) to MreB (1 µM) monitored by fluorescence anisotropy over time. Curves used to determine the fluorescent anisotropy values shown in Fig 1D. Excess (1 mM) unlabeled ATP (C) or ADP (D) was added, displacing bound ATP* and ADP*, respectively. Fitting of the dissociation phase yielded the $k_{off}$ value for ATP* and ADP* release indicated in Table 1. Differences in anisotropy amplitudes between constructs (untagged versus pFAST-tagged MreB) suggest that the N-terminal tag can influence protein folding, accessibility of the binding pocket and/or protein dynamics. (E, F) Titration experiment measuring the fluorescent anisotropy of $N^6$(6-Aminohexyl)-ATP-Atto-488 (ATP*) (E) and $N^6$(6-Aminohexyl)-ADP-Atto-488 (ADP*) (F) with MreB to determine the Kd. Used to determine fluorescent anisotropy shown in Fig 1D.

Figure S3

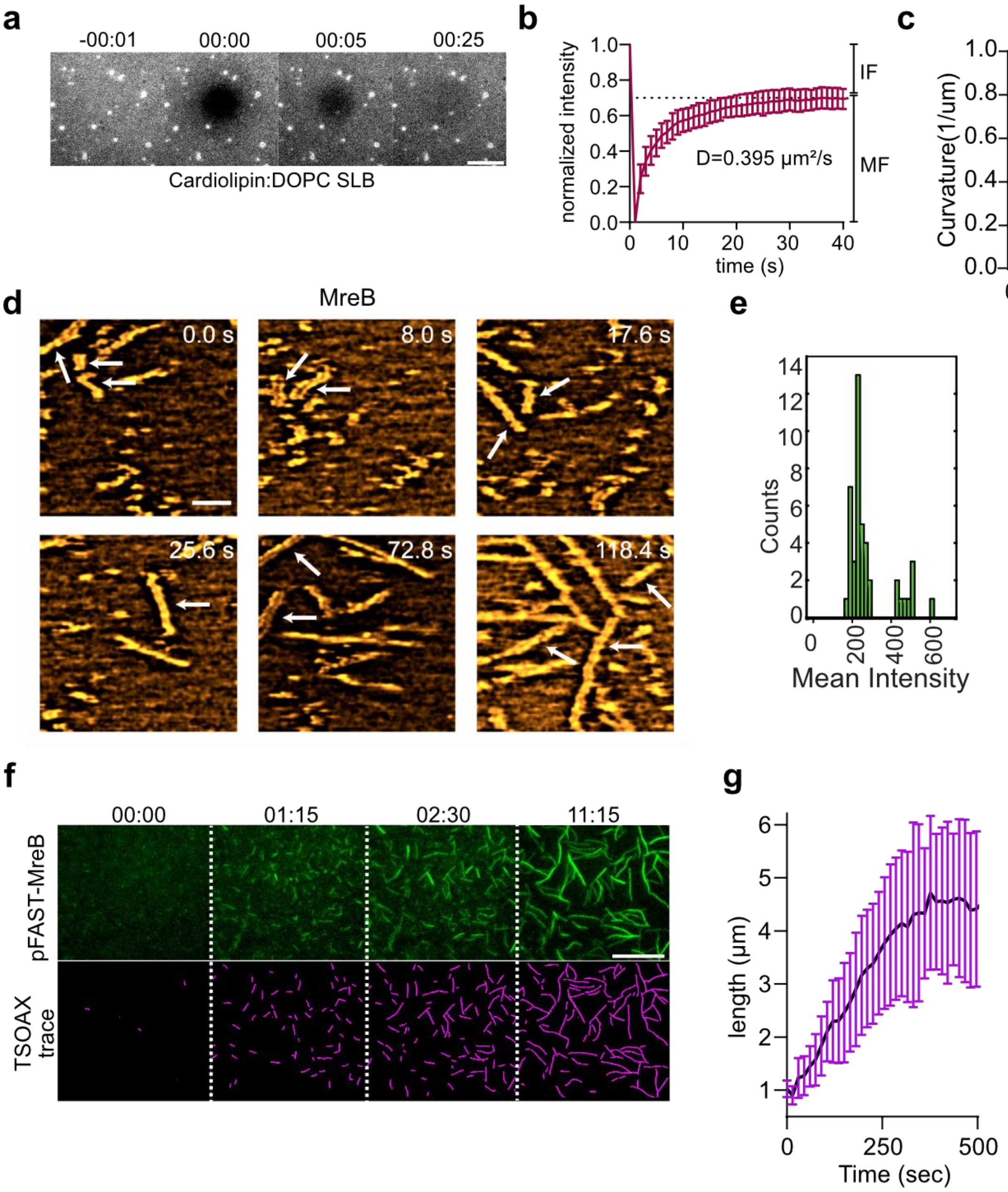

**Fig. S3. CL-containing SLBs are fluid and support pFAST-MreB polymerization.**
(A) Example of fluorescence recovery after photobleaching (FRAP) showing the fluidity of the SLBs containing 20% CL, with the remaining 80% composed of 1,2-dioleoyl-sn-glycero-3-phosphocholine (DOPC) doped with 0.2% of fluorescently labelled lipids (1,1',2,2'-tetraoleoyl cardiolipin(4-(dipyrrometheneboron difluoride)butanoyl), also known as TopFluor™ Cardiolipin). A small region of the SLB was photobleached by exposing it to a high intensity 488 nm light source, and TIRF images were acquired to monitor the evolution of the bleached spot over time. Numbers above images stand for time post photobleaching, in minutes:seconds. Scale bar, 5 µm. (B) Corresponding quantification of fluorescence intensity after photobleaching over time. Bars represent normalized mean ± standard deviation. n = 14 photobleached regions. The diffusion coefficient 'D' was determined using the halftime recovery, $0.224 \cdot w^2/t_{1/2}$=D= 0.395 µm²/s. MF,

Mobile Fraction; IF, Immobile Fraction. (C) Curvature of pFAST-MreB filaments at different densities visualized by TIRFM. Curvature was quantified using the SOAX software. ****p < 0.0001 (t test). (D) High-resolution, Fast Fourier Transform (FFT) bandpass filtered HS-AFM images showing filaments doublets (arrows) of untagged MreB (1.2 µM). Time in seconds. (E, F) Histogram of the mean fluorescence intensity along the length of filaments (n=44) (E), and time-lapse of assembly (F) of pFAST-MreB (0.05 µM) labelled with TFLime on a DOPC:CL (80:20) SLB. *Top panels*, TIRF images; *Bottom panels*, TSOAX traces used to calculate filament length over time (elongation rate). The same approach was used to determine the elongation rate of MreB with different concentrations shown in Fig. 2E. Time is indicated in minutes:seconds. Scale bar, 5 µm.

Figure S4

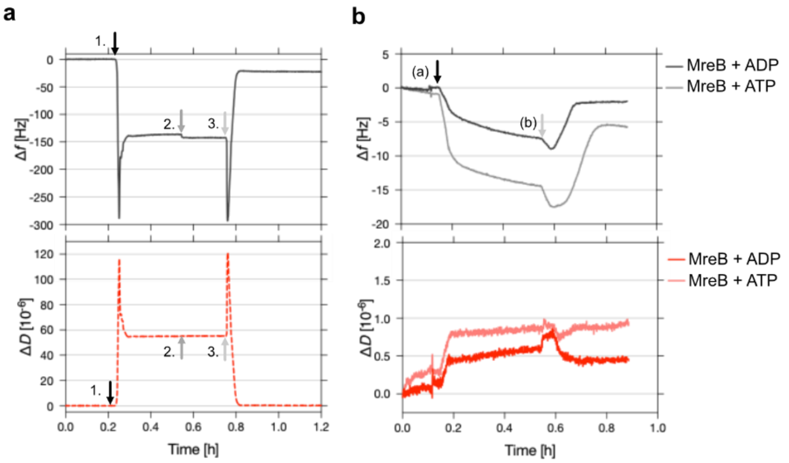

**Fig. S4. QCM-D analysis of *B. subtilis* MreB**

(A) Lipid bilayer formation on $SiO_2$ quartz crystals is represented in frequency (Δf) and dissipation (ΔD) changes over time. SLBs were formed by using solvent-assisted bilayer formation (SALB). After a stable baseline in HEPES-NaCl buffer is established, isopropanol is added (arrows #1.). Once a stable baseline in isopropanol is achieved, a lipid mixture of DOPC:CL (80:20) in isopropanol is added (#2.) starting the lipid-in-isopropanol exchange, which results in a small frequency shift. After another stable baseline, (#3.) isopropanol is exchanged with HEPES-NaCl buffer to initiate bilayer formation. (B) Subsequently, untagged MreB was (a) added to SLBs (DOPC:CL) in the presence of ATP or ADP and (b) rinsed off in MreB buffer (adsorption kinetics for 0.6 µM MreB are shown).

#### Figure S5

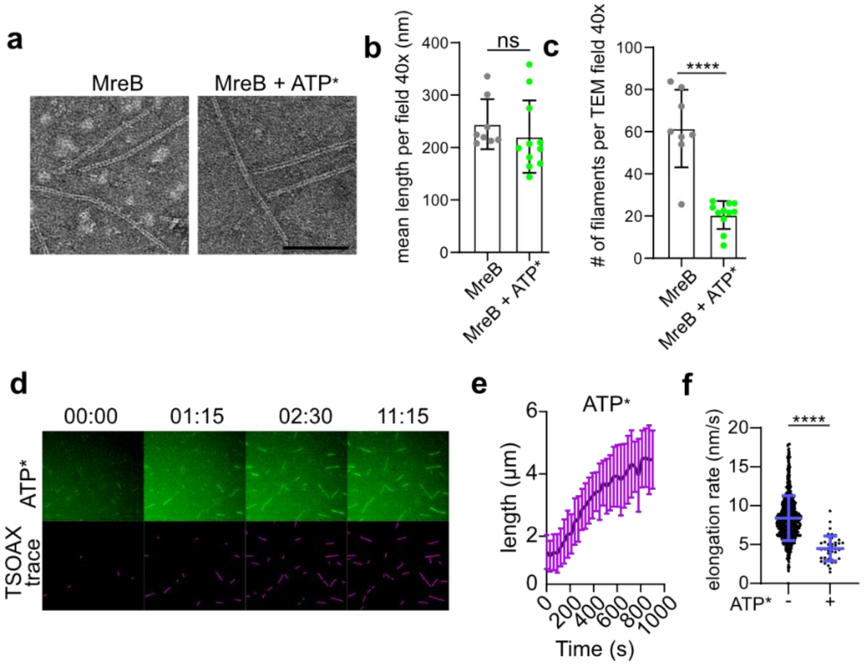

**Fig. S5. Effect of fluorescent ATP on MreB polymerization.**

(A) Representative TEM images of pairs of protofilaments formed by untagged MreB (1.5 µM) on a lipid monolayer *of B. subtilis* lipid extract in the presence of ATP (50 µM) or ATP:ATP* (50 µM : 3 µM) . Scale bar, 100 nm. (B,C) Quantification of the lengths (B) and density (C) of MreB (1.5 µM ) filaments on a lipid monolayer of *B. subtilis* total lipid extract visualized by TEM (30K magnification fields). ∗∗∗p < 0.001 (t test); ns, not significant. (D) TIRF microscopy time-lapse of polymerization of pFAST-MreB (0.05 µM ) in the presence of ATP* ($N^6$(6-Aminohexyl)-ATP-Atto-488) (ATP:ATP* 50:3 µM) on an SLB (DOPC:CL 80:20). The bottom panel shows the TSOAX traces used to calculate lengths and elongation rate. minutes:seconds. Scale bar, 5 µm. (E,F) Corresponding quantification of pFAST-MreB filament lengths (E) and elongation rates (data without ATP* also used in Fig 2E) (F).

#### Figure S6

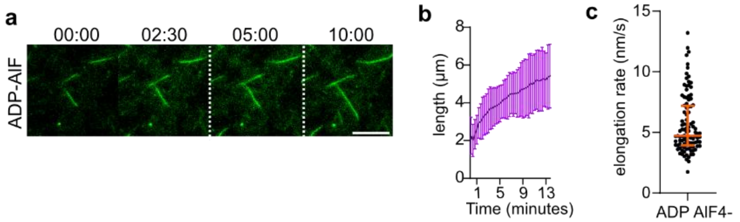

**Fig. S6. MreB filament formation in the presence of ADP-AlF4-.**

(A) Time-lapse of assembly of 0.05 µM pFAST-MreB in the presence of 0.2 mM ADP-Aluminum Fluoride (ADP-AlF4-) visualized with TFLime. Time in minutes:seconds. Scale bar, 5 µm. (B, C) Corresponding quantification of MreB filament lengths (B) and elongation rate (C).

Figure S7

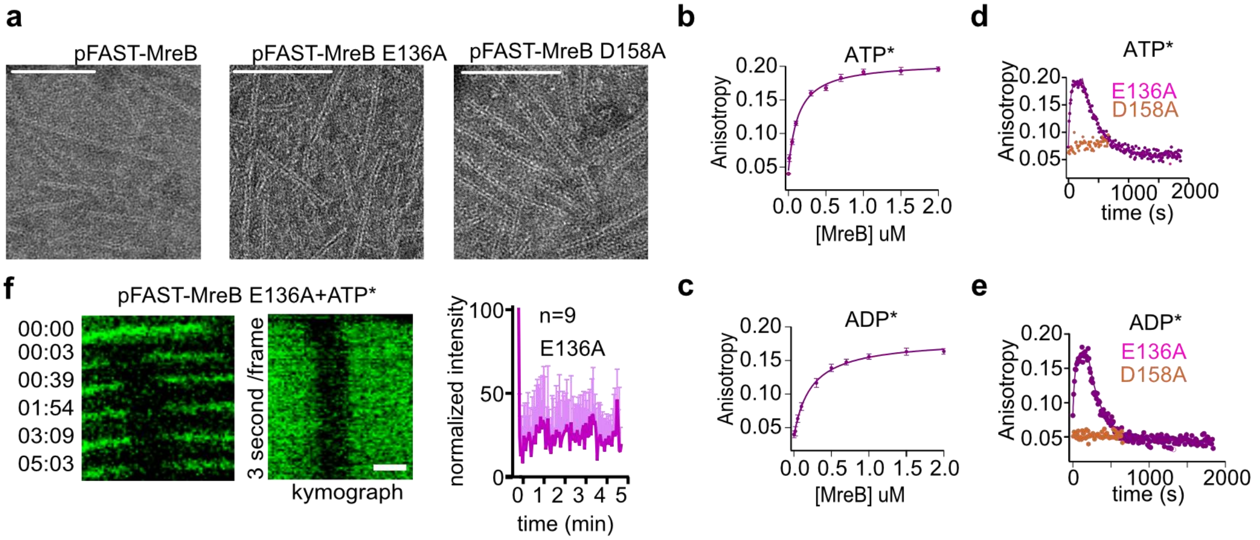

**Fig. S7. E136A and D158A filament properties.**

(A) Representative TEM images of 1.5 µM pFAST-MreB wild-type (WT), E136A and D158A polymerized on a lipid monolayer of *B. subtilis* lipid extract in the presence of ATP (2 mM). Scale bar, 100 nm. EM images show no difference between MreB WT and ATPase mutants, suggesting that nucleotide hydrolysis has no effect on MreB filament structure or mechanical properties. (B,C) Titration experiment measuring the fluorescent anisotropy of ATP* ($N^6$(6-Aminohexyl)-ATP-Atto-488) (ATP*) (B) and ($N^6$(6-Aminohexyl)-ADP-Atto-488) (ADP*) (C) with pFAST-MreB ATPase mutants for Kd determination. (D, E) Chase experiments with ATP* (D) and ADP* (E). (F) FRAP experiment on single filaments of pFAST-MreB E136A visualized with fluorescent ATP* (ATP:ATP* 17:1). *Left panel*, representative time-lapse, *middle panel*, kymograph, Time in minutes:seconds. Scale bar, 1 µm. *Right panel*, graph with FRAP result.

Figure S8

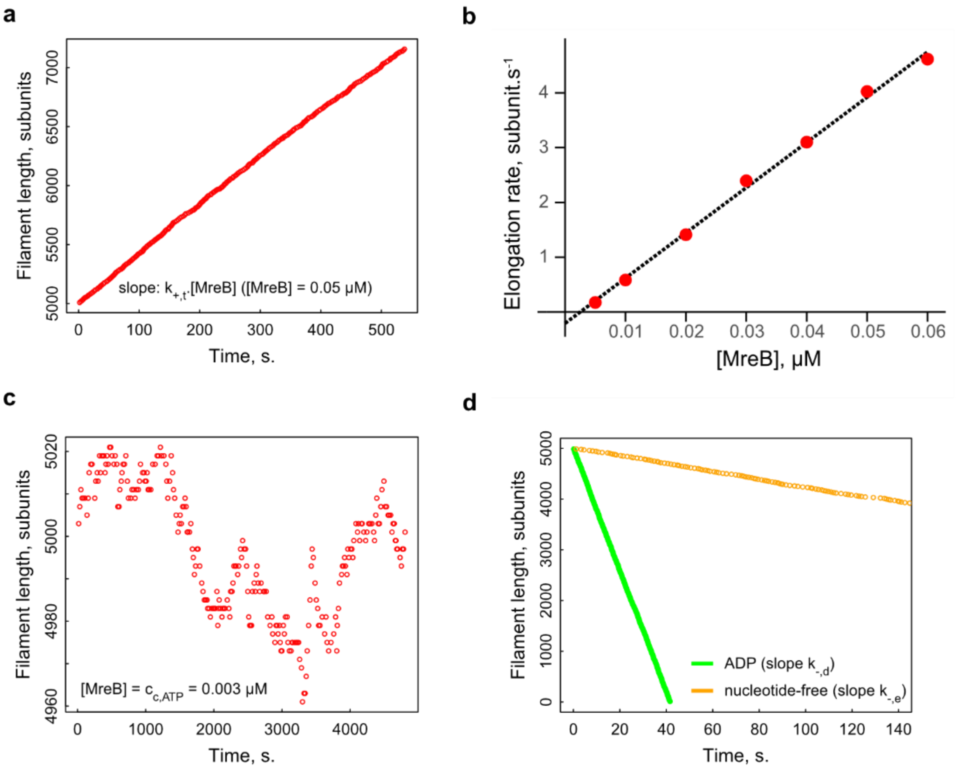

**Fig. S8. Simulating MreB filament polymerization and depolymerization.**
Rate constants used in this Figure are $k_{+,t} = 45\ \mu M^{-1}s^{-1}$, $k_{-,t} = k_{-,dp} = 0.135\ s^{-1}$, $k_{-,d} = 60\ s^{-1}$, $k_{-,e} = 4\ s^{-1}$, $k_{hyd} = k_{Pir} = 1.54 \cdot 10^{-3}s^{-1}$, and $k_{+,n} = k_{-,n} = 0$.
(A) Time evolution of MreB filament length, starting from an ATP-bound MreB filament of initial length of 5 000 subunits and a monomer concentration of [MreB-ATP] = 0.05 μM. (B) Elongation rate of MreB filaments for different MreB-ATP monomer concentrations. Intersection of linear fit (dashed line) with x axis (y = 0) indicates $c_{c,ATP} = 0.003\ \mu M$, while intersection with y axis (x = 0) indicates $k_{-,t}$. (C) Time evolution of MreB filament length, starting from an ATP-bound MreB filament of initial length of 5 000 subunits and a monomer concentration of [MreB] = $c_{c,ATP}$ = 0.003 μM. (D) Time evolution of MreB filament length, starting from a nucleotide free filament (in orange) or an ADP-bound MreB filament (in green) of initial lengths of 5 000 subunits, in the absence of MreB monomers.

#### Figure S9

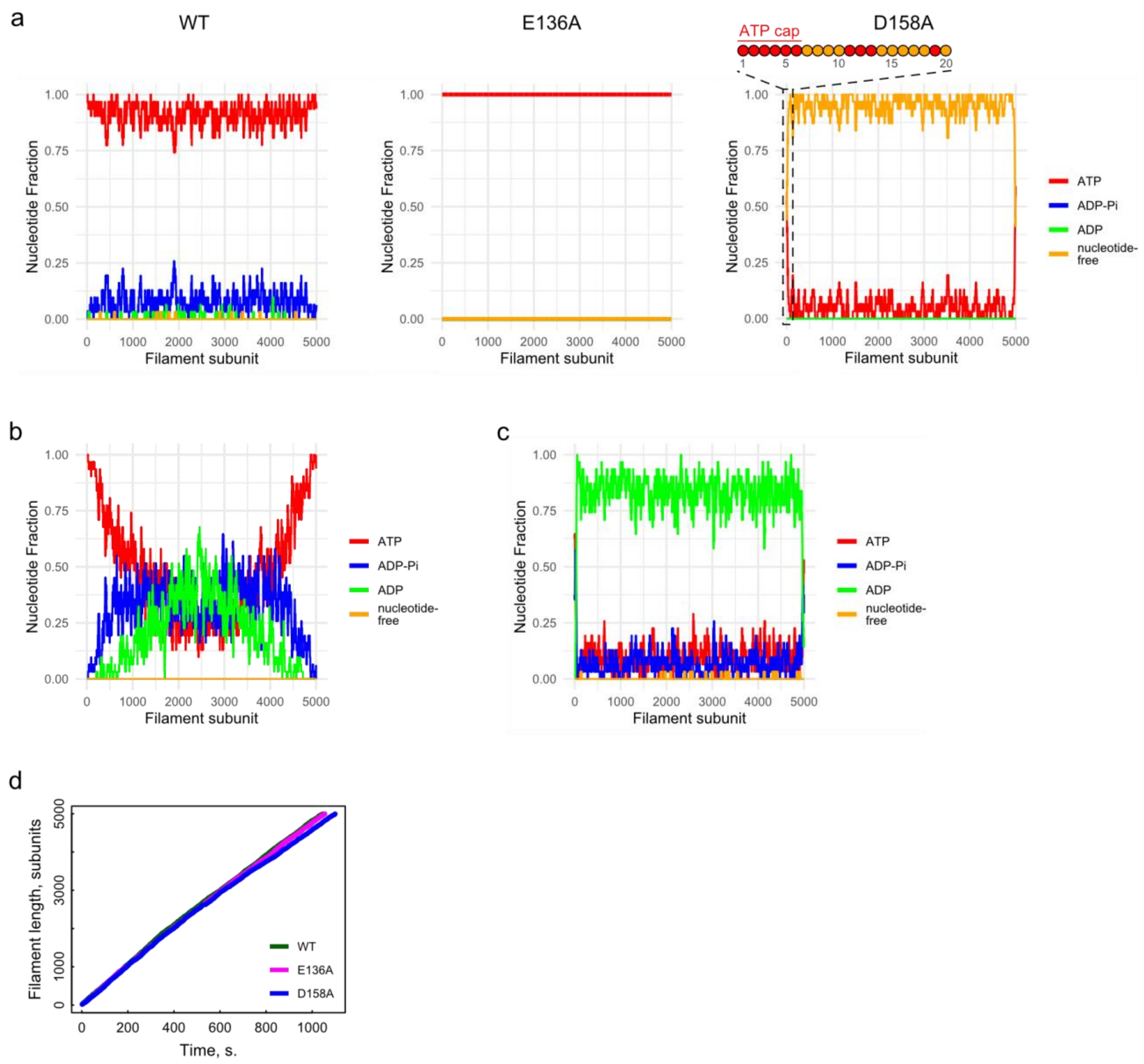

**Fig. S9. Nucleotide state of MreB filaments (WT, E136A or D158A) during polymerization.** Rate constants used in this figure include $k_{+,t} = 45\ \mu M^{-1}s^{-1}$, $k_{-,t} = k_{-,dp} = 0.135\ s^{-1}$, $k_{-,d} = 60\ s^{-1}$, $k_{-,e} = 4\ s^{-1}$.

(A) Nucleotide state of a 5000-subunit MreB filament during polymerization from MreB-ATP bound monomers, in the presence of ATP (100 µM). Left panel is a simulation of WT MreB, with $k_{hyd,WT} = k_{Pir,WT} = 1.54 \cdot 10^{-3}s^{-1}$, $k_{+,n,WT} = 0.28\ \mu M^{-1}.s^{-1}$ and $k_{-,n,WT} = 0.0141\ s^{-1}$; Central panel is a simulation of E136A MreB, with $k_{hyd,E136A} = k_{Pir,E136A} = k_{+,n,E136A} = k_{-,n,E136A} = 0$; Right panel is a simulation of D158A MreB, with $k_{hyd,D158A} = k_{Pir,D158A} = 0$. $k_{+,n,WT} = 10^{-4}\ \mu M^{-1}.s^{-1}$ and $k_{-,n,D158A} = 0.2\ s^{-1}$. Dashed box highlights the nucleotide state of the terminal subunits of the D158A filament. (B) Nucleotide state of 5000-subunit long WT MreB filament during polymerization, if intra-filament nucleotide exchange was 100-fold slower than in (A) ($k_{-,n} = 1.41\ 10^{-4}\ s^{-1}$). (C) Nucleotide state of 5000-subunit long WT MreB filament during polymerization, if ATP hydrolysis and Pi release were 100-fold slower than in (A) ($k_{hyd} = k_{Pir} = 1.54 \cdot 10^{-1}\ s^{-1}$). (D) Time evolution of WT, E136 and D158 MreB filament length in the conditions of (A).

#### Figure S9

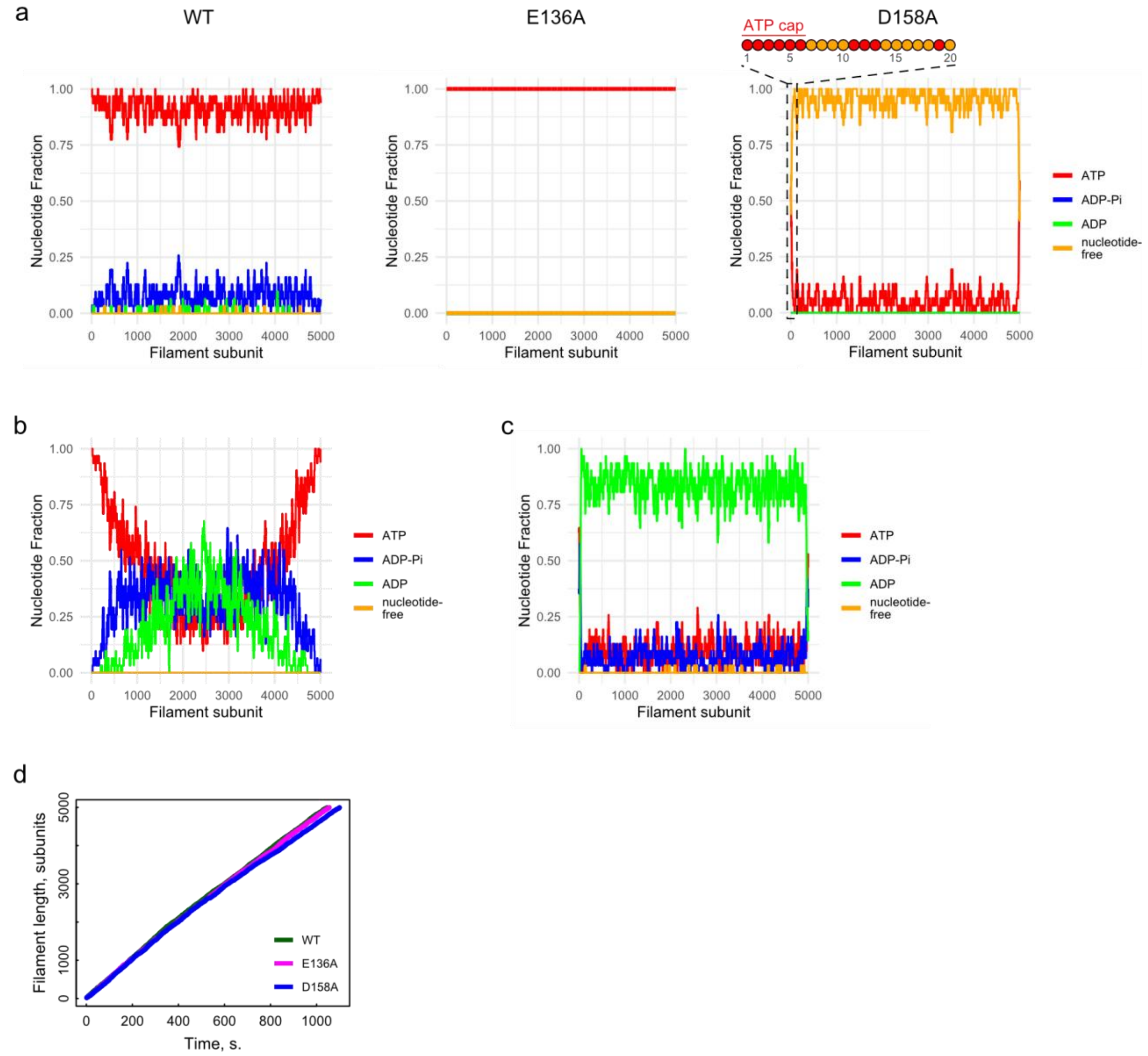

**Fig. S10. Simulation of WT, E136A and D158A MreB filaments depolymerization after flushing into a MreB-monomer free solution.**

In all panels, rate constants for WT, E136A and D158A MreB are the same as in Fig. S9A. At initial time points, the length and nucleotide state of the filaments corresponds to what they are after polymerization, as determined in Fig. S9A.

(A) Time evolution of WT MreB filament length, after flush into a nucleotide-free buffer (green curve) or into a buffer containing ADP (200 µM; magenta). (B) Time evolution of WT MreB filament nucleotide composition after flush into a nucleotide-free buffer. (C) Time evolution of WT MreB filament nucleotide composition after flush into a buffer containing ADP (200 µM; magenta). (D) Time evolution of E136A MreB filament length, after flush into a nucleotide-free buffer (green curve) or into a buffer containing ADP (200 µM; magenta). (E) Time evolution of E136A MreB filament nucleotide composition after flush into a nucleotide-free buffer. (F) Time evolution of E136A MreB filament nucleotide composition after flush into a buffer containing ADP (200 µM; magenta). (G) Time evolution of D158A MreB filament length, after flush into a nucleotide-free buffer (green curve) or into a buffer containing ADP (200 µM; magenta). (H) Time evolution of D158A MreB filament nucleotide composition after flush into a nucleotide-free buffer (F) Time evolution of D158A MreB filament nucleotide composition after flush into a buffer containing ADP (200 µM; magenta).

Figure S11

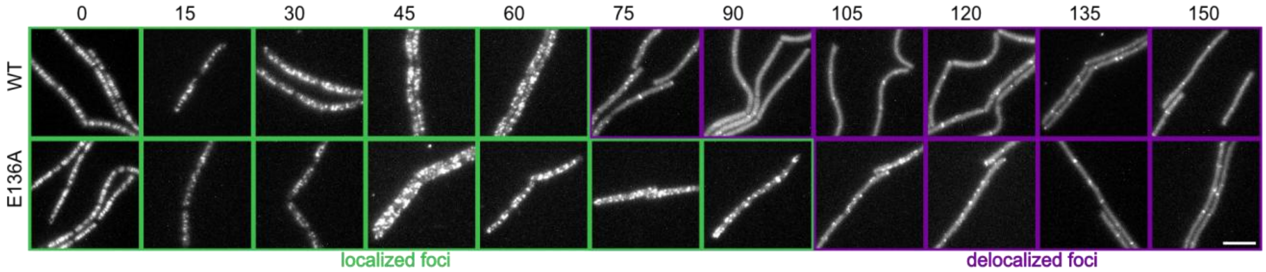

**Fig. S11. Effect of CCCP on MreB localization.**
CCCP-induced delocalization of MreB in growing cells visualized by TIRF microscopy. Cells expressing GFP-MreB WT (strain RCL421) and E136A mutant (strain RCL2028) were exposed to 100 µM CCCP. Numbers above images correspond to time post addition of CCCP (T0), in minutes. Scale bar, 5 µm.

#### Supplementary Tables S1 to S4

##### Table S1. Primers list

| Name | Sequence 5'-3' |
|---|---|
| 272-mreB | ATTGAAGCGCCTTTTGCCGCAGC |
| 273-mreB | GCAAAAGGCGCTTCAATCGGATACG |
| Ac-1116 | CATGGTTGTTGCTATCGGGGGCGGTACGACAGA |
| Ac-1117 | CCCGATAGCAACAACCATGCTTCCAGTCGGTTC |
| CC182 | GGATGTGCTCCAGTGCTTTC |
| RK14 | AATTCGAGCAGACAGACAGCCAGAAC |
| CC181 | GTCATGGGCCTTCCTATATC |
| CC272 | ATTGAAGCGCCTTTTGCCGCAGC |
| CC273 | GCAAAAGGCGCTTCAATCGGATACG |
| CC564 | TGGATCCGAAGTCTGGACATTTTAACGCGTTTGACAAACACCCA |
| CC565 | TGGGTGTTTGTCAAACGCGTTAAAATGTCCAGACTTCGGATCCA |
| CC592 | ATACATTTAAGGAGGTAAACTAATGGATTCAATAGAAAGGTAAGCATGGAACATGTTGC GTTTGG |
| CC593 | CCAAACGCAACATGTTCCATGCTTACCTTTTCTATTGAATCCATTAGTTTACCTCCTTAAAT GTAT |

##### Table S2. Strains list

| Strain | Genotype or plasmid | Construction or reference |
|---|---|---|
| ***B. subtilis*** | | |
| 168 | *trpC2* | Laboratory collection |
| RCL413 | *trpC2 Ωneo3427 ΔmreB* | [2] |
| RCL421 | *mreB*::(Kan - $P_{nat}$-rbs opt-*gfp*$^{A206K}$-*mreB*) | [28] |
| RCL925 | *mreB*::( Kan - $P_{nat}$- rbs opt-NcomGA-pFAST-*mreB*) | This manuscript |
| RCL2028 | *mreB*::(Kan - $P_{nat}$-rbs opt-*gfp*$^{A206K}$-*mreB* E136A) | This manuscript |
| ***E. coli*** | | |
| ECRCL3 | F- λ- fhuA2 [lon] ompT lacZ::T7 gene 1 gal sulA11 Δ(mcrC-mrr)114::IS10 R(mcr-73::miniTn10-TetS)2 R(zgb-210::Tn10)(TetS) endA1 [dcm] | laboratory collection |
| ECRCL281 | pET28a(+) derivative carrying 6xHIS-pFAST-MreB Bs WT under control of the T7 promoter (parental strain ECRCL3) | This manuscript |
| ECRCL623 | pET28a(+) derivative carrying 6xHIS-pFAST-MreB Bs E136A under control of the T7 promoter (parental strain ECRCL3) | This manuscript |
| ECRCL625 | pET28a(+) derivative carrying 6xHIS-pFAST-MreB Bs D158A under control of the T7 promoter (parental strain ECRCL3) | This manuscript |
| ECRCL578 | pET28a(+) derivative carrying 6xHIS-SUMO-MreB Bs WT (codon optimized for *E. coli*) under control of the T7 promoter (parental strain ECRCL3) | This manuscript |
| ECRCL592 | pET28a(+) derivative carrying 6xHIS-hsSENP1 under control of the T7 promoter(parental strain ECRCL3) | This manuscript |

Pnat = MreB natural promoter. Rbs opt = ribosome binding site optimized for *B. subtilis*. NcomGA = 8 amino acids of ComGA, which allow increased expression of the fusion protein.

**Table S3. Plasmid pET28a 6His-pFAST-MreB sequence**
**[blue – His tag, red = pFAST, green = linker, purple = *mreB*]**

TCCGGCGTAGAGGATCGAGATCTCGATCCCGCGAAATTAATACGACTCACTATAGGGGAATTGTGAGCGGATAACA
ATTCCCCTCTAGAAATAATTTTGTTTAACTTTAAGAAGGAGATATACCATGCATCATCACCATCATCACATGGAACA
TGTTGCGTTTGGAAGCGAAGATATCGAGAATACGCTTGCAAATATGGATGATGAACAACTTGACCGTTTAGCGTTT
GGTGTTATTCAACTGGATGGCGATGGGAATATCCTGCTTTATAACGCTGCAGAAGGTGACATTACAGGCAGAGATC
CGAAACAGGTCATAGGCAAGAACTTCTTCAAAGACGTAGCACCTGGAACTGATACACCCGAATTTTACGGCAAATT
CAAAGAAGGAGCAGCTTCGGGAAATCTCAACACGATGTTTGAGTGGACTATTCCGACAAGTCGTGGACCAACCAA
AGTGAAAGTACACCTGAAGAAAGCGTTGTCTGGTGACCGGTATTGGGTGTTTGTCAAACGCGTTAAAATGTCCAGA
CTTCGGATCCACGGGCCCCCCCTCGAGATGTTTGGAATTGGTGCTAGAGACCTTGGTATAGATCTTGGAACTGCGA
ATACGCTTGTTTTTGTAAAAGGAAAAGGAATTGTTGTGAGAGAGCCGTCAGTTGTCGCTTTGCAGACGGATACGAA
ATCGATTGTCGCTGTCGGAAATGATGCGAAAAATATGATTGGACGGACACCGGGCAACGTGGTGGCTCTTCGCCCG
ATGAAAGACGGCGTTATCGCTGATTATGAAACAACGGCGACGATGATGAAATATTACATCAATCAGGCCATAAAA
AATAAAGGCATGTTTGCCAGAAAACCATATGTAATGGTATGTGTCCCATCAGGCATTACAGCTGTTGAAGAACGCG
CTGTTATCGATGCGACAAGACAGGCGGGAGCGCGTGACGCGTATCCGATTGAAGAGCCTTTTGCCGCAGCAATCGG
AGCCAATCTGCCAGTTTGGGAACCGACTGGAAGCATGGTTGTTGATATCGGGGGCGGTACGACAGAAGTTGCGATT
ATTTCCCTCGGAGGCATCGTAACGTCTCAGTCAATCCGTGTAGCCGGTGATGAGATGGATGACGCGATTATCAACT
ACATCAGAAAAACGTACAATCTGATGATCGGTGACCGTACGGCTGAAGCGATTAAAATGGAAATCGGATCTGCAG
AAGCTCCTGAAGAATCCGACAACATGGAAATCCGCGGCCGCGATTTGCTCACAGGTTTGCCGAAAACAATTGAAAT
TACAGGAAAAGAGATTTCTAACGCTCTACGCGACACTGTATCTACAATTGTCGAAGCAGTGAAGAGCACACTCGAA
AAAACACCGCCTGAGCTTGCAGCAGATATCATGGACAGAGGTATAGTGTTAACCGGCGGCGGAGCGCTTTTGCGCA
ATTTGGACAAAGTCATCAGCGAAGAAACAAAAATGCCGGTCCTTATCGCCGAAGATCCGCTTGATTGTGTAGCGAT
CGGAACAGGGAAAGCACTGGAGCACATCCATCTTTTCAAAGGGAAAACTAGATAAGGATCCGAATTCGAGCTCCG
TCGACAAGCTTGCGGCCGCACTCGAGCACCACCACCACCACCACTGAGATCCGGCTGCTAACAAAGCCCGAAAGG
AAGCTGAGTTGGCTGCTGCCACCGCTGAGCAATAACTAGCATAACCCCTTGGGGCCTCTAAACGGGTCTTGAGGGG
TTTTTTGCTGAAAGGAGGAACTATATCCGGATTGGCGAATGGGACGCGCCCTGTAGCGGCGCATTAAGCGCGGCGG
GTGTGGTGGTTACGCGCAGCGTGACCGCTACACTTGCCAGCGCCCTAGCGCCCGCTCCTTTCGCTTTCTTCCCTTCCT
TTCTCGCCACGTTCGCCGGCTTTCCCCGTCAAGCTCTAAATCGGGGGCTCCCTTTAGGGTTCCGATTTAGTGCTTTAC
GGCACCTCGACCCCAAAAAACTTGATTAGGGTGATGGTTCACGTAGTGGGCCATCGCCCTGATAGACGGTTTTTCG
CCCTTTGACGTTGGAGTCCACGTTCTTTAATAGTGGACTCTTGTTCCAAACTGGAACAACACTCAACCCTATCTCGG
TCTATTCTTTTGATTTATAAGGGATTTTGCCGATTTCGGCCTATTGGTTAAAAAATGAGCTGATTTAACAAAAATTT
AACGCGAATTTTAACAAAATATTAACGTTTACAATTTCAGGTGGCACTTTTCGGGGAAATGTGCGCGGAACCCCTA
TTTGTTTATTTTTCTAAATACATTCAAATATGTATCCGCTCATGAATTAATTCTTAGAAAAACTCATCGAGCATCAA
ATGAAACTGCAATTTATTCATATCAGGATTATCAATACCATATTTTTGAAAAAGCCGTTTCTGTAATGAAGGAGAA
AACTCACCGAGGCAGTTCCATAGGATGGCAAGATCCTGGTATCGGTCTGCGATTCCGACTCGTCCAACATCAATAC
AACCTATTAATTTCCCCTCGTCAAAAATAAGGTTATCAAGTGAGAAATCACCATGAGTGACGACTGAATCCGGTGA
GAATGGCAAAAGTTTATGCATTTCTTTCCAGACTTGTTCAACAGGCCAGCCATTACGCTCGTCATCAAAATCACTCG
CATCAACCAAACCGTTATTCATTCGTGATTGCGCCTGAGCGAGACGAAATACGCGATCGCTGTTAAAAGGACAATT
ACAAACAGGAATCGAATGCAACCGGCGCAGGAACACTGCCAGCGCATCAACAATATTTTCACCTGAATCAGGATA
TTCTTCTAATACCTGGAATGCTGTTTTCCCGGGGATCGCAGTGGTGAGTAACCATGCATCATCAGGAGTACGGATA
AAATGCTTGATGGTCGGAAGAGGCATAAATTCCGTCAGCCAGTTTAGTCTGACCATCTCATCTGTAACATCATTGGC
AACGCTACCTTTGCCATGTTTCAGAAACAACTCTGGCGCATCGGGCTTCCCATACAATCGATAGATTGTCGCACCTG
ATTGCCCGACATTATCGCGAGCCCATTTATACCCATATAAATCAGCATCCATGTTGGAATTTAATCGCGGCCTAGAG
CAAGACGTTTCCCGTTGAATATGGCTCATAACACCCCTTGTATTACTGTTTATGTAAGCAGACAGTTTTATTGTTCAT
GACCAAAATCCCTTAACGTGAGTTTTCGTTCCACTGAGCGTCAGACCCCGTAGAAAAGATCAAAGGATCTTCTTGA
GATCCTTTTTTTCTGCGCGTAATCTGCTGCTTGCAAACAAAAAAACCACCGCTACCAGCGGTGGTTTGTTTGCCGGA
TCAAGAGCTACCAACTCTTTTTCCGAAGGTAACTGGCTTCAGCAGAGCGCAGATACCAAATACTGTCCTTCTAGTGT
AGCCGTAGTTAGGCCACCACTTCAAGAACTCTGTAGCACCGCCTACATACCTCGCTCTGCTAATCCTGTTACCAGTG
GCTGCTGCCAGTGGCGATAAGTCGTGTCTTACCGGGTTGGACTCAAGACGATAGTTACCGGATAAGGCGCAGCGGT
CGGGCTGAACGGGGGGTTCGTGCACACAGCCCAGCTTGGAGCGAACGACCTACACCGAACTGAGATACCTACAGC
GTGAGCTATGAGAAAGCGCCACGCTTCCCGAAGGGAGAAAGGCGGACAGGTATCCGGTAAGCGGCAGGGTCGGA
ACAGGAGAGCGCACGAGGGAGCTTCCAGGGGGAAACGCCTGGTATCTTTATAGTCCTGTCGGGTTTCGCCACCTCT
GACTTGAGCGTCGATTTTTGTGATGCTCGTCAGGGGGGCGGAGCCTATGGAAAAACGCCAGCAACGCGGCCTTTTT
ACGGTTCCTGGCCTTTTGCTGGCCTTTTGCTCACATGTTCTTTCCTGCGTTATCCCCTGATTCTGTGGATAACCGTAT
TACCGCCTTTGAGTGAGCTGATACCGCTCGCCGCAGCCGAACGACCGAGCGCAGCGAGTCAGTGAGCGAGGAAGC
GGAAGAGCGCCTGATGCGGTATTTTCTCCTTACGCATCTGTGCGGTATTTCACACCGCATATATGGTGCACTCTCAG
TACAATCTGCTCTGATGCCGCATAGTTAAGCCAGTATACACTCCGCTATCGCTACGTGACTGGGTCATGGCTGCGCC

CCGACACCCGCCAACACCCGCTGACGCGCCCTGACGGGCTTGTCTGCTCCCGGCATCCGCTTACAGACAAGCTGTG
ACCGTCTCCGGGAGCTGCATGTGTCAGAGGTTTTCACCGTCATCACCGAAACGCGCGAGGCAGCTGCGGTAAAGCT
CATCAGCGTGGTCGTGAAGCGATTCACAGATGTCTGCCTGTTCATCCGCGTCCAGCTCGTTGAGTTTCTCCAGAAGC
GTTAATGTCTGGCTTCTGATAAAGCGGGCCATGTTAAGGGCGGTTTTTTCCTGTTTGGTCACTGATGCCTCCGTGTA
AGGGGGATTTCTGTTCATGGGGGTAATGATACCGATGAAACGAGAGAGGATGCTCACGATACGGGTTACTGATGAT
GAACATGCCCGGTTACTGGAACGTTGTGAGGGTAAACAACTGGCGGTATGGATGCGGCGGGACCAGAGAAAAATC
ACTCAGGGTCAATGCCAGCGCTTCGTTAATACAGATGTAGGTGTTCCACAGGGTAGCCAGCAGCATCCTGCGATGC
AGATCCGGAACATAATGGTGCAGGGCGCTGACTTCCGCGTTTCCAGACTTTACGAAACACGGAAACCGAAGACCAT
TCATGTTGTTGCTCAGGTCGCAGACGTTTTGCAGCAGCAGTCGCTTCACGTTCGCTCGCGTATCGGTGATTCATTCT
GCTAACCAGTAAGGCAACCCCGCCAGCCTAGCCGGGTCCTCAACGACAGGAGCACGATCATGCGCACCCGTGGGG
CCGCCATGCCGGCGATAATGGCCTGCTTCTCGCCGAAACGTTTGGTGGCGGGACCAGTGACGAAGGCTTGAGCGAG
GGCGTGCAAGATTCCGAATACCGCAAGCGACAGGCCGATCATCGTCGCGCTCCAGCGAAAGCGGTCCTCGCCGAA
AATGACCCAGAGCGCTGCCGGCACCTGTCCTACGAGTTGCATGATAAAGAAGACAGTCATAAGTGCGGCGACGAT
AGTCATGCCCCGCGCCCACCGGAAGGAGCTGACTGGGTTGAAGGCTCTCAAGGGCATCGGTCGAGATCCCGGTGCC
TAATGAGTGAGCTAACTTACATTAATTGCGTTGCGCTCACTGCCCGCTTTCCAGTCGGGAAACCTGTCGTGCCAGCT
GCATTAATGAATCGGCCAACGCGCGGGGAGAGGCGGTTTGCGTATTGGGCGCCAGGGTGGTTTTTCTTTTCACCAG
TGAGACGGGCAACAGCTGATTGCCCTTCACCGCCTGGCCCTGAGAGAGTTGCAGCAAGCGGTCCACGCTGGTTTGC
CCCAGCAGGCGAAAATCCTGTTTGATGGTGGTTAACGGCGGGATATAACATGAGCTGTCTTCGGTATCGTCGTATC
CCACTACCGAGATATCCGCACCAACGCGCAGCCCGGACTCGGTAATGGCGCGCATTGCGCCCAGCGCCATCTGATC
GTTGGCAACCAGCATCGCAGTGGGAACGATGCCCTCATTCAGCATTTGCATGGTTTGTTGAAAACCGGACATGGCA
CTCCAGTCGCCTTCCCGTTCCGCTATCGGCTGAATTTGATTGCGAGTGAGATATTTATGCCAGCCAGCCAGACGCAG
ACGCGCCGAGACAGAACTTAATGGGCCCGCTAACAGCGCGATTTGCTGGTGACCCAATGCGACCAGATGCTCCACG
CCCAGTCGCGTACCGTCTTCATGGGAGAAAATAATACTGTTGATGGGTGTCTGGTCAGAGACATCAAGAAATAACG
CCGGAACATTAGTGCAGGCAGCTTCCACAGCAATGGCATCCTGGTCATCCAGCGGATAGTTAATGATCAGCCCACT
GACGCGTTGCGCGAGAAGATTGTGCACCGCCGCTTTACAGGCTTCGACGCCGCTTCGTTCTACCATCGACACCACC
ACGCTGGCACCCAGTTGATCGGCGCGAGATTTAATCGCCGCGACAATTTGCGACGGCGCGTGCAGGGCCAGACTGG
AGGTGGCAACGCCAATCAGCAACGACTGTTTGCCCGCCAGTTGTTGTGCCACGCGGTTGGGAATGTAATTCAGCTC
CGCCATCGCCGCTTCCACTTTTTCCCGCGTTTTCGCAGAAACGTGGCTGGCCTGGTTCACCACGCGGGAAACGGTCT
GATAAGAGACACCGGCATACTCTGCGACATCGTATAACGTTACTGGTTTCACATTCACCACCCTGAATTGACTCTCT
TCCGGGCGCTATCATGCCATACCGCGAAAGGTTTTGCGCCATTCGATGGTGTCCGGGATCTCGACGCTCTCCCTTAT
GCGACTCCTGCATTAGGAAGCAGCCCAGTAGTAGGTTGAGGCCGTTGAGCACCGCCGCCGCAAGGAATGGTGCAT
GCAAGGAGATGGCGCCCAACAGTCCCCCGGCCACGGGGCCTGCCACCATACCCACGCCGAAACAAGCGCTCATGA
GCCCGAAGTGGCGAGCCCGATCTTCCCCATCGGTGATGTCGGCGATATAGGCGCCAGCAACCGCACCTGTGGCGCC
GGTGATGCCGGCCACGATGCG

**Table S4. Liposome compositions**

| Name | composition | Used for |
|---|---|---|
| *B. subtilis* extract | 100 % | TEM + liposomes binding assay |
| DOPC | 100 % | Liposomes binding assay |
| DOPC + PG | 80% + 20% | Liposomes binding assay |
| DOPC + CL | 80% + 20% | Liposomes binding assay + single filament imaging (TIRF and HS-AFM) |
| DOPC + CL + PE rhodamine | 79.8% + 20% + 0.2% | Single filament imaging (TIRF and HS-AFM) |
| DOPC + CL + TopFluor™ CL | 79.8% + 20% + 0.2% | Fluidity of SLBs (FRAP) |

#### Movies captions

**Movie 1.** TIRF microscopy movie of FRAP experiment on a DOPC:CL (80%:20%) SLB visualized by TopFluor™ Cardiolipin. Time interval is 1.16 second.

**Movie 2.** HS-AFM movie of polymerization of 1.2 µM untagged MreB on a DOPC:CL (80%:20%) SLB in the presence of 0.2 mM ATP. Time interval is 0.6 seconds. Corresponding to Figure 1I.

**Movie 3.** TIRF microscopy movie of polymerization of 0.05 µM pFAST-MreB (TFLime-labelled) on a DOPC:CL (80%:20%) SLB in the presence of 0.2 mM ATP. Time interval is 15 seconds. Corresponding to Figure 1K.

**Movie 4.** HS-AFM movie of 0.6 µM untagged MreB on a DOPC:CL (80%:20%) SLB in the presence of 0.2 mM ATP. Time interval is 0.6 seconds. Corresponding to Figure 1L and 2B.

**Movie 5.** TIRF microscopy movie of 0.05 µM pFAST-MreB (TFlime-labelled) filaments on a DOPC:CL (80%:20%) SLB in the presence of 0.2 mM ATP. Time interval is 15 seconds. Corresponding to Figure 1N, O.

**Movie 6.** TIRF microscopy movie of pFAST-MreB (0.05 µM) filaments formed in the presence of ATP after replacing the buffer with MreB-free buffer containing ADP (0.2 mM). Time interval is 15 seconds. Corresponding to Figure 4A.

**Movie 7.** HS-AFM movie of MreB (0.6 µM) filaments formed in the presence of ATP (0.2 mM) after replacing ATP in the buffer with 0.2 mM ADP. Time interval is 0.4 seconds. Corresponding to Figure 4E.

**Movie 8.** TIRF microscopy movie of TFLime-labelled pFAST-MreB E136A (0.05 µM) filaments after replacing the buffer with MreB-free buffer containing ADP (0.2 mM). Time interval is 15 seconds. Corresponding to Figure 4A.

**Movie 9.** TIRF microscopy movie of pFAST-MreB WT filaments capped with E136A filament caps, visualized by fluorescent ATP (ATP*), after replacing the buffer with MreB-free buffer containing ADP (0.2 mM). Time interval is 20 seconds. Corresponding to Figure 4F, G, H.
